# Rapid canalisation of mandible structure in Tetrapoda

**DOI:** 10.1101/2025.03.17.643812

**Authors:** Emily C. Watt, Ryan N. Felice, Anjali Goswami

## Abstract

Evolution of the mandibular structure has played a key role in the taxonomic diversification and niche exploration of tetrapods, from the initial water-to-land transition to the origin and radiation of mammals. While jaw evolution through these key transition points have long attracted attention, no studies of jaw evolution to date capture the breadth of tetrapod diversity and evolutionary history. Here, we reconstruct the compositional evolution of the tetrapod mandible with a comprehensive dataset of 36 traits scored in 3047 species. This sampling spans the earliest eotetrapodiforms to extant amphibians, mammals, and reptiles and birds, capturing immense ecological, developmental, and taxonomic diversity among a near-even split of extant and fossil tetrapods. We find that lower jaw morphology rapidly canalises at the base of each major clade, including, Eotetrapodiformes, Amniota, Synapsida, Sauropsida, and Amphibia, as well as subclades within those groups, and that, as a result, jaw compositional variation is highly phylogenetically structured. We estimate an average of over 100 shifts in three key traits: number of elements (dermal and endochondral bones), number of tooth-bearing elements, and total teeth per hemimandible, with symmetrical rates of gains and losses estimated for the number of elements, and correlated evolution between these three characters. Although novel elements arise occasionally, disparity in jaw composition overwhelmingly decreases as clades canalise rapidly, successively, and pervasively through tetrapod evolutionary history.

## Main text

The water-to-land transition was a pivotal moment in vertebrate evolution (Clack, 2012; Ahlberg, 2018), heralding the expansion of the eotetrapodiforms (Coates & Friedman, 2010), whose descendants became the exceptionally successful and diverse Tetrapoda, the superclass of limbed vertebrates that includes amphibians, mammals, and birds and reptiles. Tetrapods have diversified significantly since those initial excursions onto land, expanding onto all continents, traversing earth, air, and water, and adapting to suit a huge variety of ecological niches (e.g., Sahney *et al*., 2010; Saladin *et al*., 2019; Maher *et al*., 2022; Saban *et al*., 2023). Shifting from an obligate aquatic lifestyle to land ∼390 million years ago radically modified the morphology and behaviour of the earliest eotetrapodiforms, including alterations to their limbs and locomotion (e.g., Coates *et al*., 2002; Shubin *et al*., 2006; Clack, 2009; Pierce *et al*., 2012; Standen *et al*., 2014; Amaral & Schneider, 2018) and respiratory systems (e.g., Randall, 1981; Coates & Clack, 1991; Brainerd, 1999, 2015; Clack, 2007; Cupello *et al*., 2022). Another region of considerable modification is the feeding system, specifically the lower jaw (Figure 1), reflecting the shifts from feeding in the water to processing food and eating on land (e.g., Olson, 1961; Taylor, 1987; De Vree & Gans, 1994; Anderson *et al*., 2013). Subsequently, the tetrapod lower jaw as a structure has undergone a wide variety of adaptations to suit a diverse array of feeding styles, including lingual feeding (such as in frogs), inertial feeding (such as in varanid lizards), and suction feeding (such as in turtles) (Montuelle *et al*., 2009; Wainwright *et al*., 2015; Bramble & Wake, 1985).

**Figure 1:**
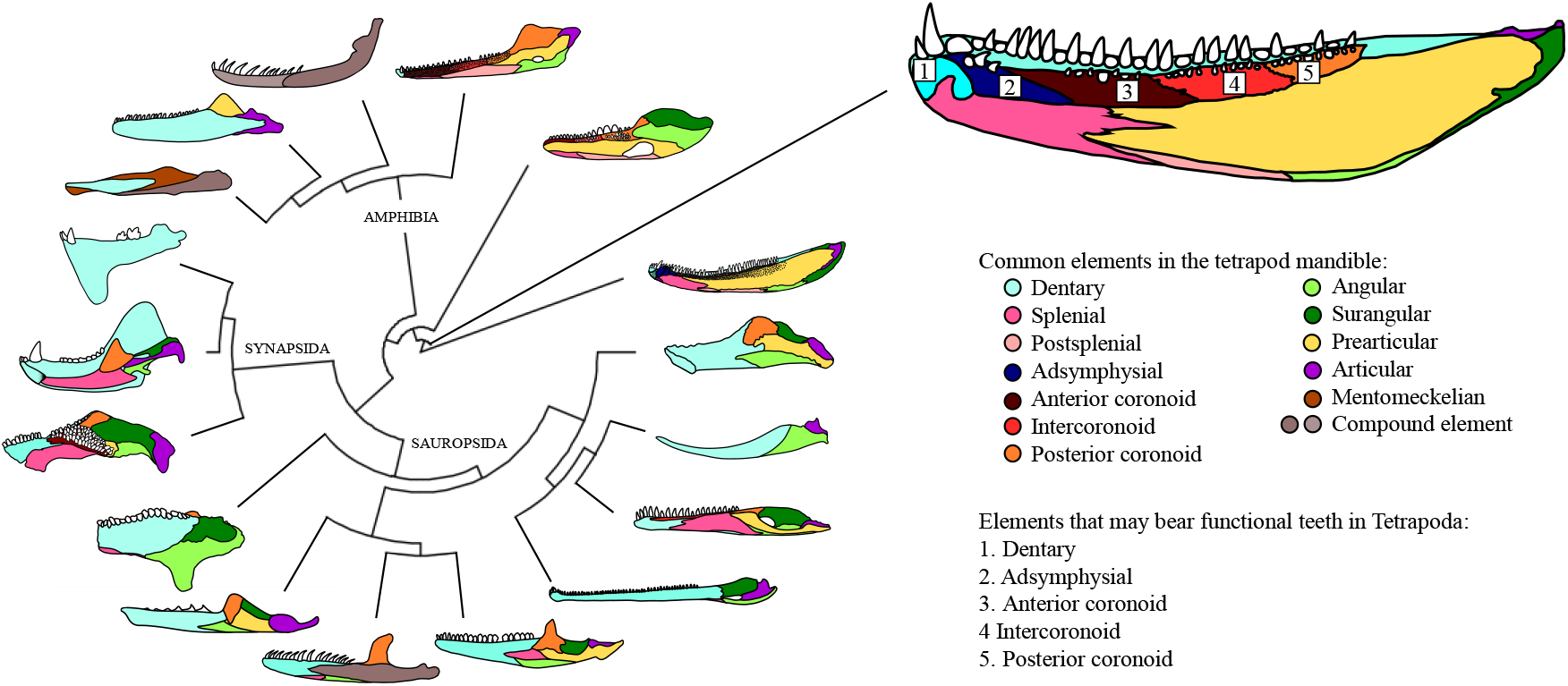
Select variation in the composition and morphology of the tetrapod lower jaw. Colours denote the most common elements in the jaw, and numbered are the elements that can bear functional teeth.

Whereas the changes in structure, morphology, and feeding mechanisms across Tetrapoda since the water-to-land transition are well known, lower jaw composition has not been comprehensively studied and compared across the whole clade. Indeed, while the evolution of the lower jaw of the earliest tetrapods was the focus of a classic study (Ahlberg & Clack, 1998), and much attention has been paid to the transitioning of jaw elements to the middle ear in mammals (e.g., Crompton, 1963a, 1963b; Luo *et al*., 2016, 2017; Navarro-Díaz *et al*., 2019), no studies have considered the shifts in jaw composition across the whole of Tetrapoda. As a result, there is no evidence to understand whether the lower jaw is canalised (robust against environmental or genetic influences (Debat & David, 2001; Hallgrimsson *et al*., 2019)) early in Tetrapoda, or whether the evolution of the tetrapod lower jaw has been persistently shaped by innovation and novelty.

Accordingly, here we reconstruct shifts in lower jaw composition over nearly 400 million years of evolutionary history by collating 36 discrete traits across 3047 tetrapod species, spanning the taxonomic breadth of Tetrapoda. Importantly, we pair a dataset containing a large extinct component (53.5%) with a modern phylogenetic framework, sampled at least to family level, which captures the vast history of evolutionary change through time, including direct data on the polarity and timing of morphological shifts. Specifically, we aim to: (i) quantify the disparity of jaw composition through Tetrapoda, (ii) determine whether jaw complexity, as captured by number of elements, decreases through time; and finally, (iii) quantify whether losses occur more frequently than gains for three key traits (number of elements, teeth, and tooth-bearing elements [see Methods for details] in the hemimandible).

## Results

### Morphospace occupation

Compositional variation of the jaw is striking on two accounts. The first is that stem tetrapods occupy a much greater range of variation than the more derived tetrapods (Figure 2). Interestingly, this higher volume of morphospace occupation is representative of not just the earliest Devonian and Carboniferous eotetrapodiforms and polyphyletic lepospondyls, but also includes stem taxa for the two major tetrapod clades: the temnospondyls (stem Amphibia) and the stem amniotes (including anthracosaurs, chroniosuchians, diadectomorphs, embolomeres, and seymouriamorphs).

**Figure 2:**
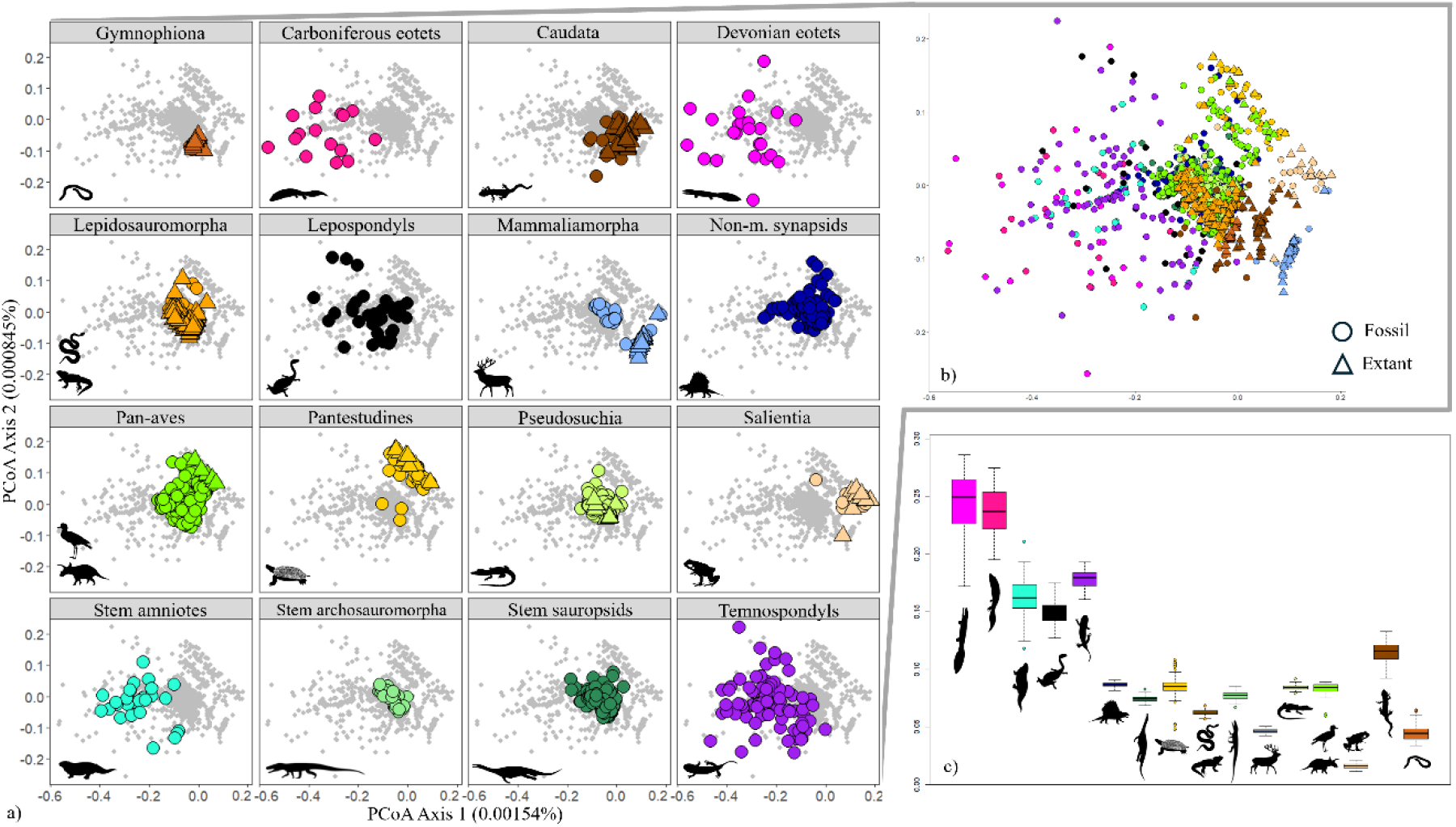
(a) Principal coordinate analysis (PCoA) of all tetrapod jaw characters across individual clades (coloured), and (b) across the entire dataset for PCo axes 1 and 2. Extant species indicated with triangles; extinct species indicated with circles. (c) Estimated disparity of each group using PCOA output with bootstrapping (n=100). Vectors used here and in subsequent figures are from Phylopic.org under CC1.0 Universal Public Domain Dedication, Attribution 3.0 Unported, Attribution-NonCommercial 3.0 Unported, and Attribution 4.0 International licenses: Oophaga pubilio (Salientia) by Chuanxin Yu, Triturus marmoratus (Caudata) by Beth Reinke, Dermophis mexicanus (Gymnophiona) and Crocodylus moreletii (Pseudosuchia) by Jose Carlos Arenas-Monroy, Glyptemys insculpta (Pantestudines) by Gabriela Palomo-Munoz, Crotalus viridis (Serpentes) by Blair Perry, Burhinus oedicnemus (Aves) by Auckland Museum, Triceratops horridus (Dinosauromorpha) by Matt Dempsey, Cervus elaphus (Mammalia) by Ferran Sayol, Sphenodon punctatus (Rhynchocephalia) by Steven Traver, Euparkeria capensis (Stem archosauromorpha) by Oliver Demuth, Diadectes (stem amniotes) by Dmitry Bogdanov (original drawing) and Roberto Díaz Sibaja (vectorisation), Dimetrodon (Synapsida) by Alessio Ciaffi, Nothosaurus mirabilis (stem sauropsids) and Loxomma allmanni (Carboniferous eotetrapodiforms) by Dan Niel, Diplocaulus (Lepospondyli) by Gareth Monger, and Balanerpeton woodi (Temnospondyli)and Acanthostega gunnari (Devonian eotetrapodiforms) by Scott Harman.

Secondly, much of the morphospace occupation among the more derived tetrapods is phylogenetically structured. This is most apparent among clades with extant species, particularly each lissamphibian order and Mammaliamorpha. The salientians, for example, have the lowest volume of morphospace occupation across Tetrapoda (Figure 2c); however, the region of morphospace that they occupy is invaded by few other taxa. Yet, other extant clades such as Lepidosauromorpha and Pseudosuchia display little variation outside of the general pattern set early in Amniota.

We first examine morphospace occupation by clade, and then by time. PCo1 explains nearly twice the variation of PCo2 and three times that of PCo3 (0.00154% vs. 0.000845% vs. 0.000491 [3 s.f.] respectively; SI).

### Morphospace occupation by clade

Morphospace occupation is highly disparate in the earliest tetrapods, particularly the Devonian and Carboniferous eotetrapodiforms, but also the stem amniotes, temnospondyls, and polyphyletic lepospondyls. Together, these early taxa occupy the full range of variation explained by PCo2, and occupy the negative extreme of PCo1, a range of variation unmatched by any of the derived clades. In contrast to the disparity of the stem amniotes, the non-mammaliamorph synapsids and stem sauropsids each occupy more restricted regions of the morphospace and display the more generalised amniote jaw composition. This centralised region of morphospace is also occupied by stem archosauromorphs, pseudosuchians, and the majority of both Pan-aves and Lepidosauromorpha.

As a distance-based method, individual PCo axes cannot be distilled into a few influential traits; however, the distribution of clades provides insight into the characters driving the major patterns of variation across Tetrapoda. For instance, in the aforementioned central region of jaw morphospace, the clustered taxa overwhelmingly share six or seven jaw elements, with little variation between the elements present (dentary, splenial, angular, surangular, prearticular, articular, and one coronoid element). The region defined by positive extreme of PCo1 and negative extreme of PCo2 is largely occupied by Mammaliamorpha, a clade characterised by unique reduction of jaw elements, ultimately down to a single bone. The overlap between some mammaliamorphs and salientians at the positive extreme of PCo1, however, suggests that this axis is not entirely driven by the number of elements in the jaw and is influenced by other characters, such as edentulism (most salientians and some mammaliamorphs: e.g. monotremes and derived mysticetes). Additionally, the region defined by positive extremes of both PCo1 and PCo2 shows two diagonal clusters of taxa that both lack dentition in the jaw, but the taxa within the upper cluster also bear beaks. For instance, the anomodonts, a clade of non-cynodont therapsids which has toothed, edentulous, and edentulous and beaked individuals, are neatly divided between these two clusters, with all toothed anomodonts falling into the central generalised region of the morphospace.

The axes that explain less variation correspond more with autapomorphies of a single clade. For example, PCo3 captures variation of the number of teeth in the jaw in the caecilians, whose jaw compositions are largely identical, except in the numbers of teeth and of tooth-bearing elements in their jaws. The caecilians fall neatly into a line that closely corresponds to the number of teeth, from the oldest fossil specimen, which has the most teeth and falls on the positive extreme of PCo3, while caecilians with fewer teeth fall at the negative end of this axis (Figure S1). However, this pattern does not follow for any other clade on PCo3, which is notable particularly as caecilians themselves do not have the highest number of teeth of any tetrapod.

### Morphospace occupation by time

Analysing the pattern of morphospace occupation through time (Figure 3) demonstrates a shift from diverse jaw compositions in the Devonian to narrower ranges of jaw composition through time. Certainly, by the Permian, the pattern of the generalised amniote jaw composition is realised by the earliest stem sauropsids and non-mammaliamorph synapsids. Clades with greater novelty in jaw composition than the general amniote pattern emerge by the Jurassic, such as Lissamphibia, Mammaliamorpha, and Pantestudines. Each of these clades occupy a distinct region of the morphospace, invaded by few other taxa. While disparity in morphospace occupation initially declines through the Palaeozoic, the variance broadly remains comparable from the Permian onwards (Figure 3b). We find no significant relationship between the variance of successive time periods, and therefore also no impact of the major mass extinction events on morphological variation.

**Figure 3:**
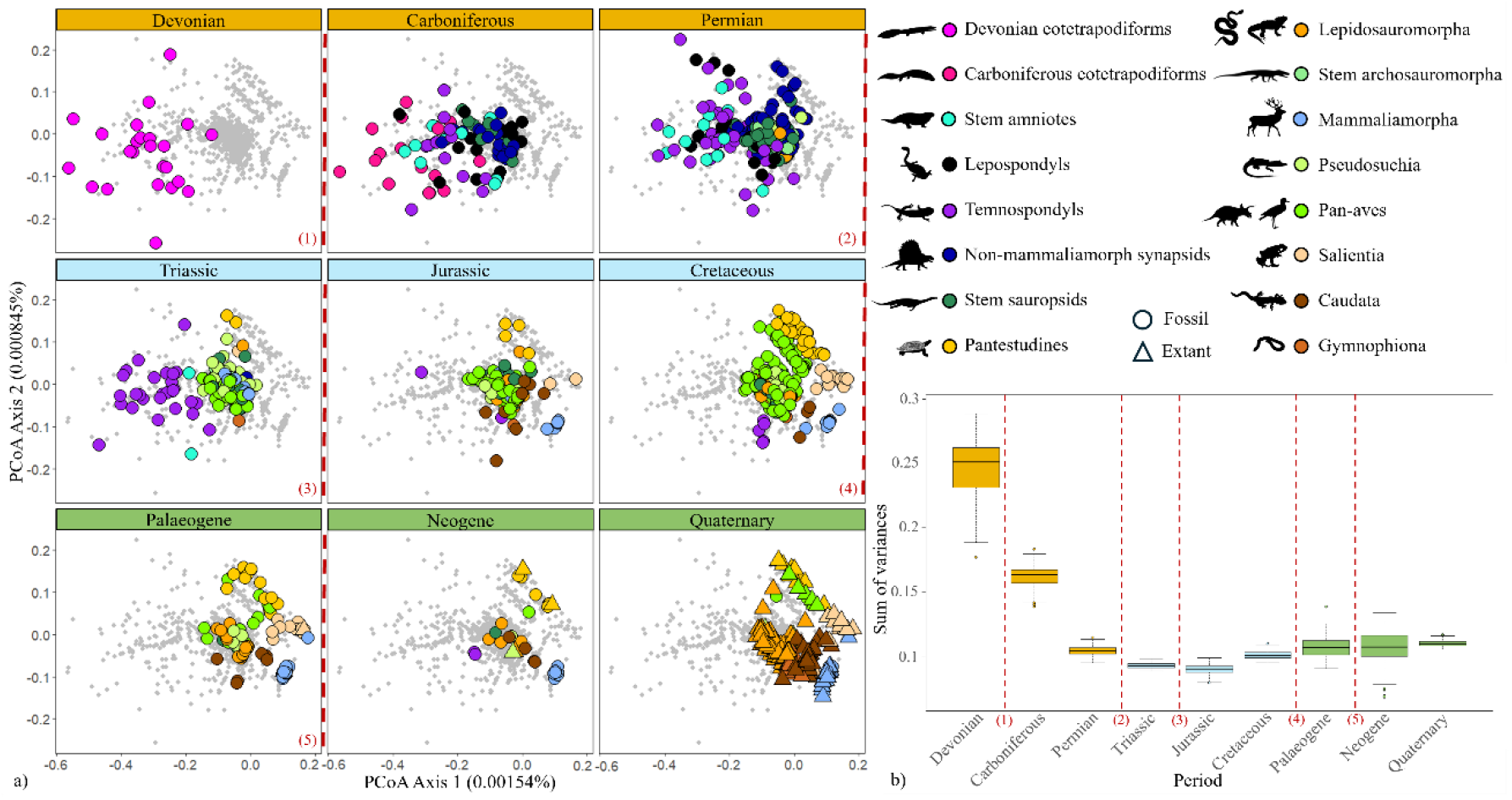
(a) Split PCoA of all tetrapod jaw characters, with each graph indicating the disparity binned by geological period (British Geological Survey, 2020), and (b) sum of variances for each time bin. Tip data was used to separate species into time periods. Red dashed lines indicate major mass extinction events: 1) End Devonian, 2) Permo-Triassic, 3) Triassic-Jurassic, 4) Cretaceous-Palaeogene (K-Pg), and 5) Eocene-Oligocene.

### Trends in tetrapod jaw composition

The number of total elements in the jaw is highest at the base of the tree, with the earliest tetrapod jaws consisting of eleven elements (or twelve - *Eusthenopteron foordi*, Porro *et al*., 2015) (Figure 4). Consistently through all major clades, the stem taxa show the most variation and the higher numbers of jaw elements, and by the extant taxa, the numbers of jaw elements have largely reached a stasis. While the highest numbers of jaw elements are mostly restricted to the base of the tree, the albanerpetontid temnospondyls, which persist to the Pliocene and are possibly nested within Lissamphibia (Matsumoto & Evans, 2018), retain ten elements in their jaws, as in most other temnospondyls (except *Cacops aspidephorus* and *Plagiosuchus pustuliferus* which have eleven elements, Damiani *et al*., 2009; Anderson *et al*., 2020). Higher numbers of elements are also gained in parts of Tetrapoda, for instance, in the non-avian dinosauromorphs, in which, several taxa gain novel elements (predentary, antarticular), and others gain/regain a second coronoid element. Overwhelmingly, however, the number of elements in the jaw has decreased through time, and the major clades have canalised very early in their evolutionary histories. Most extant clades have little variation in the number of jaw elements among taxa, with the most variation found in Caudata. A small amount of variation is also found in Lepidosauromorpha (Dibamidae bear five elements, a minority of taxa within Lepidosauromorpha bear six elements, and most bear seven), and likewise Pantestudines variably bearing either five (only *Dermochelys coriacea* (Evers *et al*., 2022)), six, or seven elements. Similar patterns of early canalisation of the major clades, and overall decreasing complexity through time are also seen in the numbers of tooth-bearing elements and overall numbers of teeth in the hemimandible (Figures S2 and S3).

**Figure 4:**
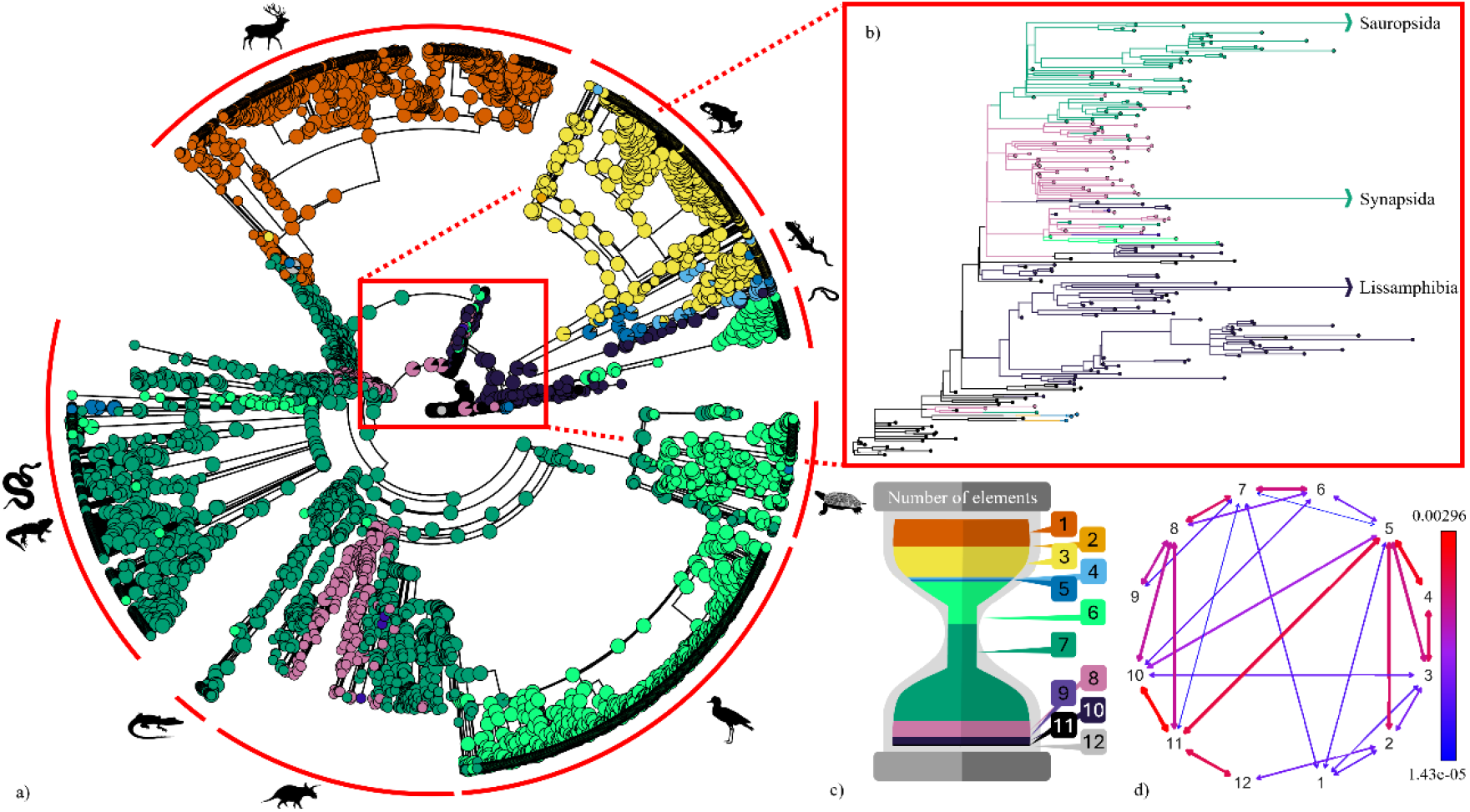
(a) Stochastic character map of the number of overall elements in the hemimandible under a symmetric model of trait evolution. (b) Inset box shows close-up of tetrapod stem, including Devonian and Carboniferous early tetrapods, paraphyletic lepospondyls, temnospondyls barring albanerpetontids (possibly nested within Lissamphibia (Matsumoto & Evans, 2018)), and stem amniotes, including stem sauropsids and stem non-mammaliamorph synapsids. Tip colours reflect observed character states, internal node colours and branch colours on the inset reflect mapped ancestral state estimates. (c) The proportion each state was estimated to occupy across the tree, and (d) a rate transition matrix showing transitions estimated to occur more frequently in red and less frequently in blue.

Evolutionary modelling of the numbers of teeth, tooth-bearing elements, and elements in the hemimandible estimates different rates for gains and losses for the numbers of teeth and tooth-bearing elements, and symmetrical rates for gains and losses for the number of elements (Table S1). Across the 100 simulations for the number of elements under a symmetrical model, we found an average of 144.84 estimated changes between states, with cumulative increases ranging between 36-141 and cumulative decreases between 38-138. Although bearing seven elements was found to be the most commonly estimated and observed state (Figure 4a), faster transitions were found with five, eight, and eleven elements (Figure 4b). For the number of teeth in the hemimandible, the highest transition rate is from a high number of teeth to a moderate number of teeth, with the next highest rate the inverse, although this was 30 times less frequent (0.06 and 0.002 respectively) (Figure S3). In all cases, the transition from a higher number of tooth-bearing elements to a lower number (loss) is faster than the inverse (gain), except between four and five tooth-bearing elements, for which gains are marginally faster than losses (Figure S2). Additionally, we find that support for correlated evolution of each pair of the three characters: numbers of teeth, tooth-bearing elements, and elements in the hemimandible (Table S2). As with the models of evolutionary rates, we find that the correlated models between elements and another state are symmetrical, whereas between numbers of teeth and tooth-bearing elements, correlated all-rates-differ is best supported.

## Discussion

We find here that canalisation is also a pervasive theme in the evolution of jaw composition in Tetrapoda. Under canalisation, there is an expectation of a relationship between the observed characters and estimated disparity, with an expected wider morphospace occupation of more ancestral species, as in early burst evolutionary models (e.g., Hughes *et al*., 2013), and a more constricted morphospace occupation of descendants. These results support this longstanding evolutionary hypothesis that the majority of variation will be found at the base of the clade, with declining variation amongst more derived taxa. We find this to hold not only for Tetrapoda, but for Amphibia and Amniota respectively, and for each major clade within. We estimate the ancestral jaw at the root of the tetrapod clade as bearing eleven elements, including five tooth-bearing elements, and a moderate (between one and 50) number of teeth. As expected under a model of canalisation, we identify that each subsequent clade has reduced the number of elements, tooth-bearing elements, and teeth in the jaw. Canalisation in this data therefore represents the narrowing of available evolutionary options in the structure of the lower jaw, with innovation in terms of jaw composition generally involving the reduction of elements.

Whereas canalisation can be easy to conceptualise as an ever narrowing set of options that can no longer be expanded, novelty can nonetheless occur within canalisation (Debat & David, 2001; Palmer, 2004). For example, although Sauropsida displays widespread similarities in overall jaw composition, it is nonetheless the main location of novelty in jaw composition. The predentary of ornithischian dinosaurs is the most common novel element in Sauropsida. This ossification has been suggested to be similar to the most widespread novel element across Tetrapoda (Ferigolo & Langer, 2007): the mentomeckelian, an ossification of cartilage at the anterior of the jaw found in most lissamphibians (e.g. Late Jurassic stem-caudatan *Lingolongtriton daxishanensis*, Jia & Gao, 2019), except the salamander families Plethodontidae, Sirenidae, and Amphiumidae (Carroll, 2007). Comparable structures to the mentomeckelian have also been identified in some extant lizards and birds (Ferigolo & Langer, 2007) and the captorhinid *Moradisaurus grandis* (Modesto *et al*., 2018), as well as outside of Tetrapoda, in some jawed fishes (mentomandibular: Grande & Bemis, 1998). The presence of the mentomeckelian clusters *Moradisaurus* with the caecilians on PCo3 and PCo4, although other taxa in the dataset with mentomeckelians (some caudatans and salientians) do not cluster with this grouping.

Other novel elements in Sauropsida include the antarticular, a small dermal element situated close to the articular near the posterior of the mandible, which has been identified in three genera of theropod dinosaurs, *Asfaltovenator* (Rauhut & Pol, 2019), *Allosaurus* (Madsen, 1976; Sasso *et al*., 2018; Chure & Loewen, 2020), and *Bagaraartan* (Osmólska, 1996), and the supradentary, found among some theropod and sauropodomorph dinosaurs and non-avian maniraptoriforms and considered be homologous with the intercoronoid/middle coronoid of earlier tetrapods (e.g., Brown & Schlaikjer, 1940; Hurum & Currie, 2000; Brochu, 2003).

In concert with the gains of novel elements across Tetrapoda, we find that gains in elements across the data are as likely as losses. While novelty explains a proportion of this trend, many of the novel elements such as the mentomeckelian, predentary, and to a lesser extent, the supradentary, are relatively pervasive across whole subclades. In contrast, taxa in other clades, such as Pantestudines, continually switch between the number of elements in their jaw, in a less phylogenetically structured manner, likely driving the symmetry of gains and losses across Tetrapoda, rather than the novel elements themselves.

Another exception to the pervasive canalisation in tetrapod jaw composition is found within Lissamphibia. Despite similar early canalisation, lissamphibians and amniotes demonstrate different ranges of compositional variation. On the whole, all lissamphibian orders have differing morphospace occupations than each other, which is in contrast to the majority of Sauropsida, whose subclades largely occupy the centralised region of the morphospace, and share common features in their jaw compositions as noted previously. While Salientia and Gymnophiona comprise the lowest and second lowest volumes of morphospace occupation of all higher level clades, the regions they occupy are mostly unique. Both orders clearly demonstrate the effects of rapid and extreme canalisation of jaw composition. In contrast, Caudata occupies a particularly large region of morphospace, following the earliest tetrapod clades (Devonian and Carboniferous eotetrapodiforms, temnospondyls, lepospondyls, and stem amniotes) and near double that of any other extant clade (0.0204; the next highest extant clade is Pantestudines: 0.0108). Caudata are unique among tetrapods for having incredibly complex and varied life histories, in particular, many taxa are paedomorphic, retaining larval characteristics throughout their lifespans (Rose, 2003). The retention of larval characteristics among some caudatans is likely driving the variation in this data, resulting in a broader morphospace occupation, higher disparity, and less extreme canalisation of jaw composition relative to other extant tetrapod clades.

Jaw evolution in crown tetrapods has been conceptualised as being constrained by neuromuscular “software” (Liem, 1985), with feeding ‘hardware’ (jaws, skull, muscles, and dentition) modified for feeding specialisations (Bramble & Wake, 1985). Trait decoupling is one evolutionary pathway by which morphological and functional specialisations have been hypothesised to occur within constrained systems, and has been identified between the oral and pharyngeal jaws of cichlid fish (Hulsey *et al*., 2006; Conith & Albertson, 2021), between hearing and mastication of the mammalian lower jaw (Tseng *et al*., 2023), and between the morphology and biomechanical disparity of the lower jaw in herbivorous dinosaurs (MacLaren *et al*., 2017). Interestingly, we find that although dentition, or lack thereof, contributes strongly to the disparity in tetrapod jaw composition, dentition and jaw elements are best explained by correlated evolutionary models. We find that the signal relating to the number of elements in the jaw is particularly strong, weighting the AICc scores of the correlated symmetrical models as much more significant than the other models.

It is perhaps surprising that to recover no decoupling of the number of teeth and elements in the jaw, given that the jaw of Mammalia, which constitutes nearly 6.5% of the dataset, is characterised by broadly only one element but complex dentition. However, considering Synapsida as a whole, we identify the same persistent canalisation of jaw composition through time which is identified as a common theme throughout each major clade (Figure 5). The earliest synapsids occupy the highest volume of the morphospace in comparison to more derived taxa; double that of the non-cynodont therapsids (0.0123 vs. 0.0062, respectively). Interestingly, the Permian non-cynodont therapsids show an exploration of jaw composition into that occupied largely by panaves and pantestudines. This is driven by the a subset of anomodonts, heavy-set herbivores with particularly robust jaws specialised for propalinal sliding and that are edentulous or have osseous “beaks” (see Benoit *et al*., 2018). Morphospace occupations is progressively restricted from non-mammaliamorph cynodonts onwards, with an abrupt shift away from the more generalised amniote bauplan of non-mammaliamorph synapsids to an otherwise unoccupied morphospace region by crown Mammalia. While the evolution of the synapsid lower jaw has been an extensive focus of research and is often held as a unique case within tetrapods (e.g., Sidor, 2003; Luo *et al*., 2015; Navarro-Díaz *et al*., 2019; Morales-García *et al*., 2021), we find that it is actually a representative system which reflects the overall pattern of canalisation across Tetrapoda.

**Figure 5:**
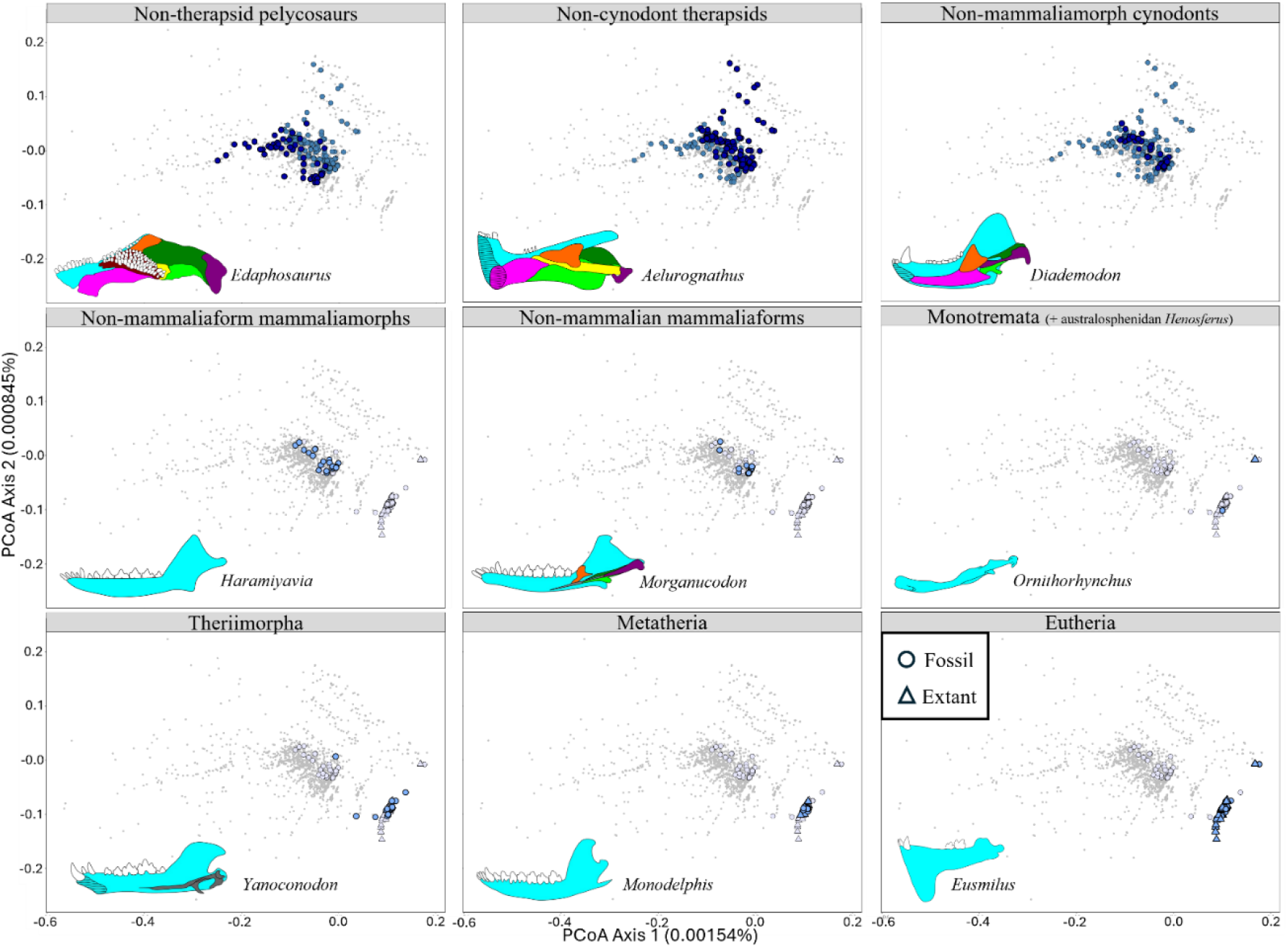
Principal coordinate analysis (PCoA) of all tetrapod jaw characters across the entire dataset (grey), with the non-mammaliamorph synapsids (dark blue) and Mammaliamorpha (light blue) indicated by larger points for PCo axes 1 and 2. Extant species indicated with triangles, extinct species indicated with circles. Examples of jaws from each group have been given as schematic diagrams, with individual jaw elements coloured per Figure 1.

Overall, we identify decreasing complexity and disparity of jaw composition from the earliest tetrapods to the extant clades, with the largest shifts occurring in the late Palaeozoic. Jaw composition canalised rapidly at the emergence of Tetrapoda, at the base of each major clade, and the base of each successive subclade. Morphospace occupation is strongly phylogenetically structured among Salientia, Gymnophiona, and Mammaliamorpha, indicating extremes of canalisation, whereas the majority of sauropsids retain a more ancestral amniote bauplan. While we identify a clear pattern of decreasing elements through time, there are numerous gains, including the pervasive lissamphibian mentomeckelian and ornithischian predentary. Gains are not always as static, and this state switching of the number of elements explains the estimation of support for a symmetrical evolutionary model. Overwhelmingly, diversity of jaw composition is the greatest at the base of clades, with the same pattern repeating at each clade level: rapid canalisation.

## Materials and Methods

### Sampling and character collation

We scored 36 discrete and meristic lower jaw characters across 1629 fossil and 1418 extant tetrapods (n=3047), including 30 stem amniotes, 714 synapsids (303 extant), 179 stem sauropsids, 154 pantestudines (80 extant), 40 stem archosauromorphs, 874 pan-avians (including dinosaurs, with 450 extant birds), 127 pseudosuchians (17 extant), 362 lepidosauromorphs (233 extant), and 397 lissamphibians (335 extant: 46 caecilians, 61 salamanders and newts, 228 frogs and toads). We sampled to family level where possible for extant species, and attempted to record every described basal fossil jaw per clade regardless of level of completeness. Where data was only recorded for a genus, we implemented synonyms, which are usually the type species for that genus, but may also be older binomial names in order to match with the composite phylogeny. Many extant birds were scored from the matrix of (Livezey & Zusi, 2006), which recorded data predominantly per genus, but also occasionally to family level. Here we also implemented synonyms, including the type species and up to four other species per genus; where a genus has over five species, we capped the recording at five so as not to overinflate the data. We treated data from the matrix ascribed to family level similarly, identifying the genera in that family and recorded a representative number of up to five species.

We scored characters based mainly on published literature, with some data on extant squamates and birds scored from high-resolution micro-CT scans (SI). For each species, the characters were scored for only one side (hemimandible). The characters scored include the presence or absence of elements within the jaw and tooth counts for each dentigerous element and for the hemimandible overall. Where elements were partially or fully fused, they were recorded as their individual components. Where characters were not preserved they were marked as missing, with the exception of total number of elements. For incomplete fossils, the total number of elements in the jaws were inferred to have no more elements than their closest relatives based on phylogenetic bracketing, as indicated in SI. Where data was provided in the literature, we also recorded heterodonty/homodonty, presence or absence of mandibular fenestrae, presence or absence of a beak, tooth implantation type, state of symphyseal fusion, and presence of Meckel’s cartilage separate to elements formed from Meckelian ossification (e.g. articular).

In the tooth counts, we included anything described as a tooth from the main tooth row, marginal teeth, accessory teeth, fangs, and fang-like teeth. We excluded denticles, as these are often undescribed, or the total number is unknown due to the preservation of the specimen. In at least one stem amniote, *Eoherpeton watsoni*, the prearticular element bears a field of denticles (Smithson, 1985). As we excluded denticles in the overall tooth count, we also excluded the prearticular from counts of the number of tooth-bearing elements. Additionally, we excluded pseudoteeth from the tooth counts; these are usually projections of dermal bone of the dentary in small projections that appear tooth-like, and have been identified in birds, turtles, and squamates. In this instance, if pseudoteeth were present and no true teeth were present on the dentary, the dentary tooth count would be scored as 0, and therefore, where applicable, overall tooth count and overall tooth presence would also be scored as 0. We initially discretised the tooth count data using the function ‘Mclust’ from the R package ‘mclust’ (Scrucca *et al*., 2023) in the R computing environment version 4.4.1 (R Core Team, 2024), however we found that the numbers of clusters did not attenuate, so discriminating the optimal number of categories left uncertainty, even using the Bayesian information criterion (BIC). While these clusters represent the density of tooth counts in the data, we found them less biologically meaningful. We therefore used three clusters, representing zero teeth (edentulism), a moderate number of teeth (1-50 teeth) and a high number of teeth (51+ teeth). In particular, we felt that this better represented the high quantity of edentulous jaws in the dataset than the density-driven ‘mclust’ clusters (Figure S4).

We created a composite phylogeny using Mesquite 3.61 (Maddison & Maddison, 2021) and the R packages ‘ape’ (Paradis & Schliep, 2019), ‘geiger’ (Pennell *et al*., 2014), and ‘phytools’ (Revell, 2012). We created a fossil backbone spanning Tetrapoda, using an initial backbone phylogeny (Ruta *et al*., 2018), which focussed on the earliest tetrapods and amphibians. We also created individual composite trees for early amniotes and sauropsids, synapsids, and pantestudines, which were independently dated and then grafted together, and inserted into the original early tetrapod tree. For each tree, we started with a fossil backbone phylogeny (pantestudines, Foth & Joyce, 2016; early amniotes and sauropsid tree, Ford & Benson, 2020; non-mammalian synapsids, Hellert *et al*., 2023). Placeholder tips for extant clades were inserted using Mesquite. We estimated divergence times and branch lengths using the function ‘bin_cal3TimePaleoPhy’ from the R package ‘paleotree’ (Bapst, 2012). The extant clades were then taken from recent molecular analyses (Aves: Jetz *et al*., 2012; Squamata: Tonini *et al*., 2016; Mammalia: Upham *et al*., 2019) and pasted onto the placeholder branches using getMRCA() from ‘ape’ and paste.tree() from ‘phytools’. We combined the early amniotes and sauropsids, synapsids, and pantestudines trees using these functions and retained and rescaled the dated branch lengths through the packages ‘dispRity’ (Guillerme, 2018) and ‘RRphylo’ (Castiglione *et al*., 2018). We followed the Archelosauria hypothesis for the placement of pantestudines, which has been supported by genomic data (e.g., Crawford *et al*., 2015; Lyson & Bever, 2020) and some modern morphological hypotheses (e.g. Simões *et al*., 2022).

We consider that the polyphyletic and taxonomically challenging lepospondyls are an important facet of early tetrapod evolution, in spite of the inherent challenges in including them in analyses, and thus have retained their presence herein.

### Variation and disparity in jaw composition through time

To identify the major axes of variation in jaw composition, we conducted a Principal Coordinates analysis (PCoA) on all 36 characters for the full dataset of 3047 species. We used the ‘vegdist’ function with the method ‘gower’ from the R package ‘vegan’ (Oksanen *et al*., 2020) and used the ‘pcoa’ function from the R package ‘ape’. We applied Lingoes correction for negative eigenvalues, which returned the relative weightings of each axis of the PCoA. We assessed the disparity in PCoA scores between clades using the package ‘dispRity’. We used the ‘dispRity.per.group’ function to estimate disparity for each group with bootstrapping (n=100), and the ‘test.dispRity’ function with the method ‘gower’ and ‘sequential’ comparisons to perform a permutational MANOVA (‘adonis.dispRity’) to assess significance between the disparity of successive time periods.

### Ancestral states and transition rates of key traits

To reconstruct changes in key jaw traits across the phylogeny, we selected the three traits that were best represented in the data: number of elements, number of tooth-bearing elements, and number of teeth in the hemimandible. The number of overall teeth had three states (0 teeth, 1-50 teeth, 51+ teeth), the number of elements had twelve states (between 1 and 12 elements), and the number of dentigerous elements had six states (between 0 and 5 elements). We only included taxa for which the data on all three characters were available (2611 species, SI). In ‘phytools’ we used the ‘fitMk’ function to fit extended Mk models, for each of the three characters under equal rates (ER), symmetrical (SYM), and all-rates-differ (ARD). We compared the fit of the models using weighted AIC scores with the ‘aic.w’ function.

We then used the ‘make.simmap’ function from the R package ‘phytools’ to generate a distribution of 100 stochastic character maps, using an equal rates model for the number of teeth and tooth-bearing elements, and a symmetrical model for the number of elements, based on the outcome of the evolutionary modelling analyses. We estimated ancestral states for the same three key characters as above (number of elements, tooth-bearing elements, and teeth), calculating the averaged probability of the state of the internal nodes across the 100 iterations, and mapping the averaged nodes onto one random stochastic character map. We extracted the posterior probability distributions of state changes from the 100 iterations using the function ‘density’ (from base R package ‘stats’).

We also tested the correlation between the number of elements and the number of tooth-bearing elements (2611 species) using the R package ‘corHMM’ (O’Meara *et al*., 2022). corHMM uses a Markov model to describe the discrete transitions between the observed states. We ran six models, across two different model structures: correlated ER, independent ER, correlated SYM, independent SYM, correlated ARD, and independent ARD. We compared model fit using AICc scores.

## Acknowledgments

For thought-provoking and valuable discussion on modelling discrete data we thank Gustavo Burin, Natalie Cooper, Julien Clavel, and Thomas Guillerme and for suggestions on coding we are grateful to Liam Revell and Katherine Corn. For discussions on amphibian jaws, we thank the members of the NHM Herpetology Group, Susan Evans, Marc Jones, and Anne Claire-Fabre, and for discussions on early tetrapod jaws we thank Laura Porro. This work was supported by funding from London Natural Environmental Research Council DTP grant number NE/S007229/1 (to ECW) and European Research Council grant STG-2014-637171 (to AG).

## Author Contributions

Conceptualisation, ECW, RNF, and AG; Methodology, ECW, RNF, and AG ; Investigation, ECW; Formal Analysis, ECW, RNF, and AG; Writing – Original Draft, ECW; Writing – Review & Editing, ECW, RNF, and AG; Supervision, RF and AG.

