## Supplementary material for "Rapid canalisation of mandible structure in Tetrapoda": SI

The PDF file includes:

1. Table of supplementary figures ........................................................................ 1
2. Table of supplementary tables ........................................................................ 12
3. Composite tree references .............................................................................. 17

### 1. Supplementary figures

Figure S1: Principal coordinate analysis (PCoA) of all tetrapod jaw characters across (a) individual clades (coloured), and (b) across the entire dataset (grey) for PCo axes 3 and 4.

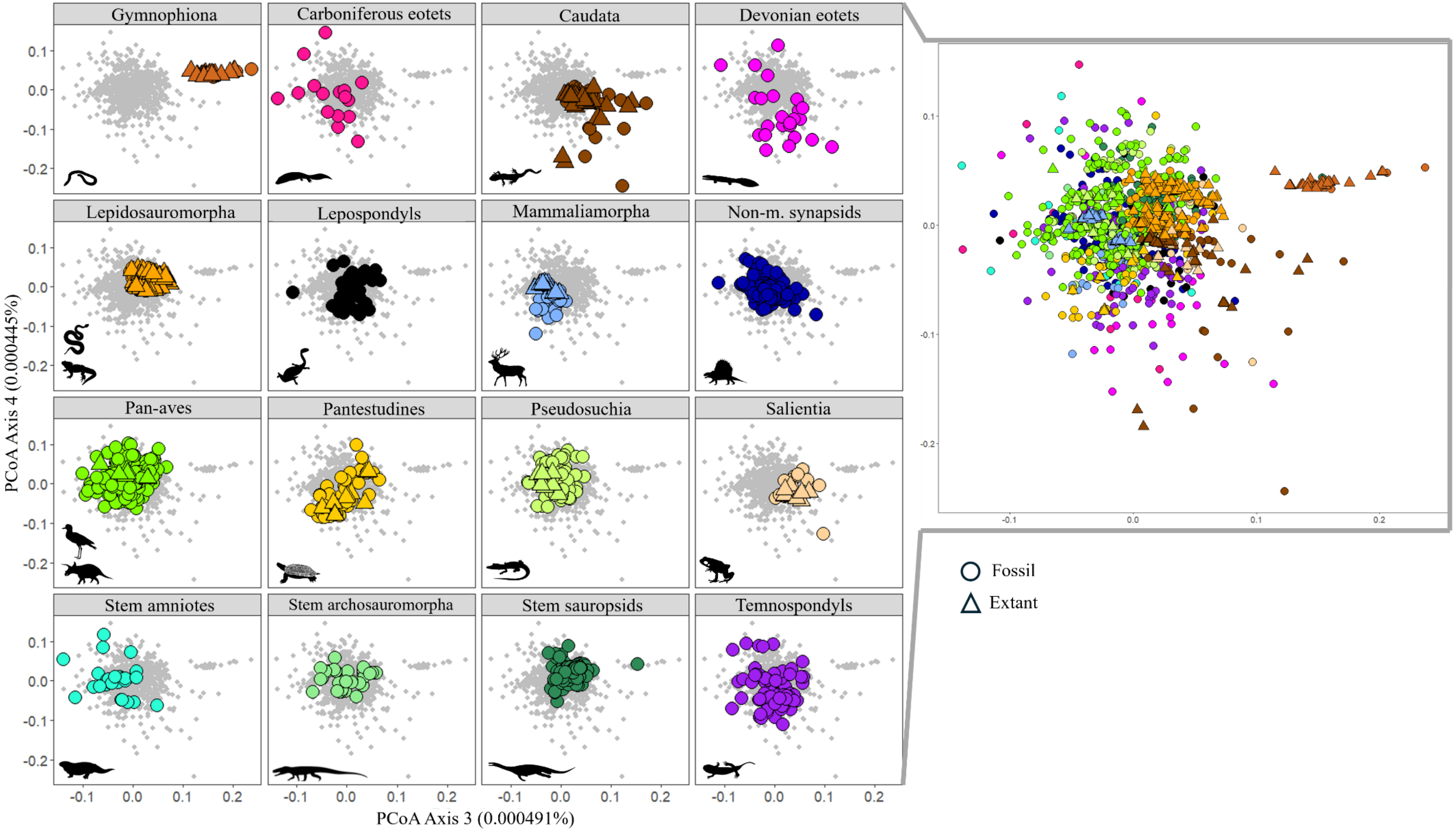

Figure S2: Stochastic character map of the number of tooth-bearing elements in the hemimandible (all-rates-differ (ARD)), across (a) the full dataset and (b) the base of the clade. Tip colours reflect observed character states, internal node colours and branch colours on the inset reflect mapped ancestral state estimates. (c) Rate transition matrix showing transitions estimated to occur more frequently in red and less frequently in blue.

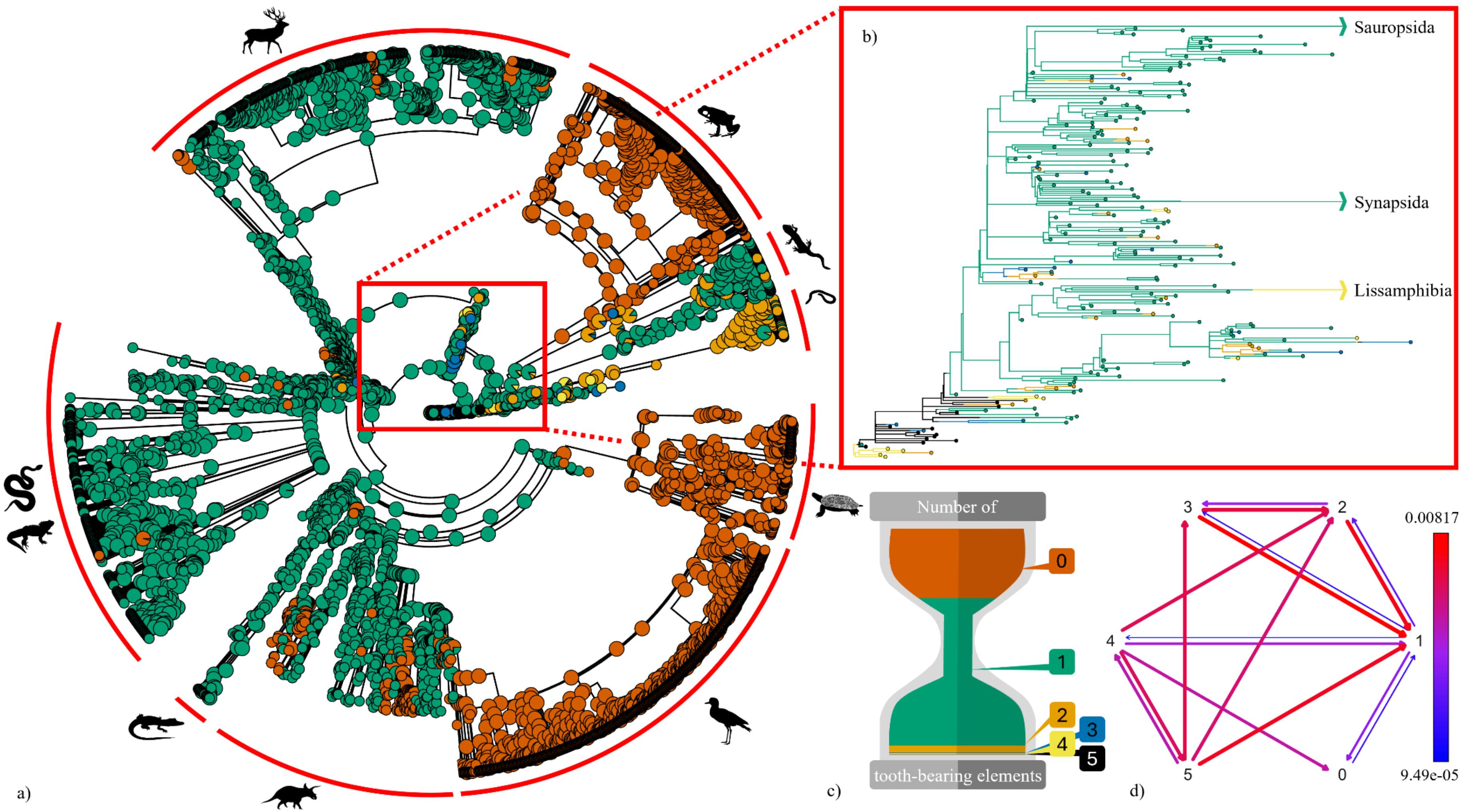

Figure S3: Stochastic character map of the number of teeth in the hemimandible (all-rates-differ (ARD)), across (a) the full dataset and (b) the base of the clade. Tip colours reflect observed character states, internal node colours and branch colours on the inset reflect mapped ancestral state estimates. (c) Rate transition matrix showing transitions estimated to occur more frequently in red and less frequently in blue.

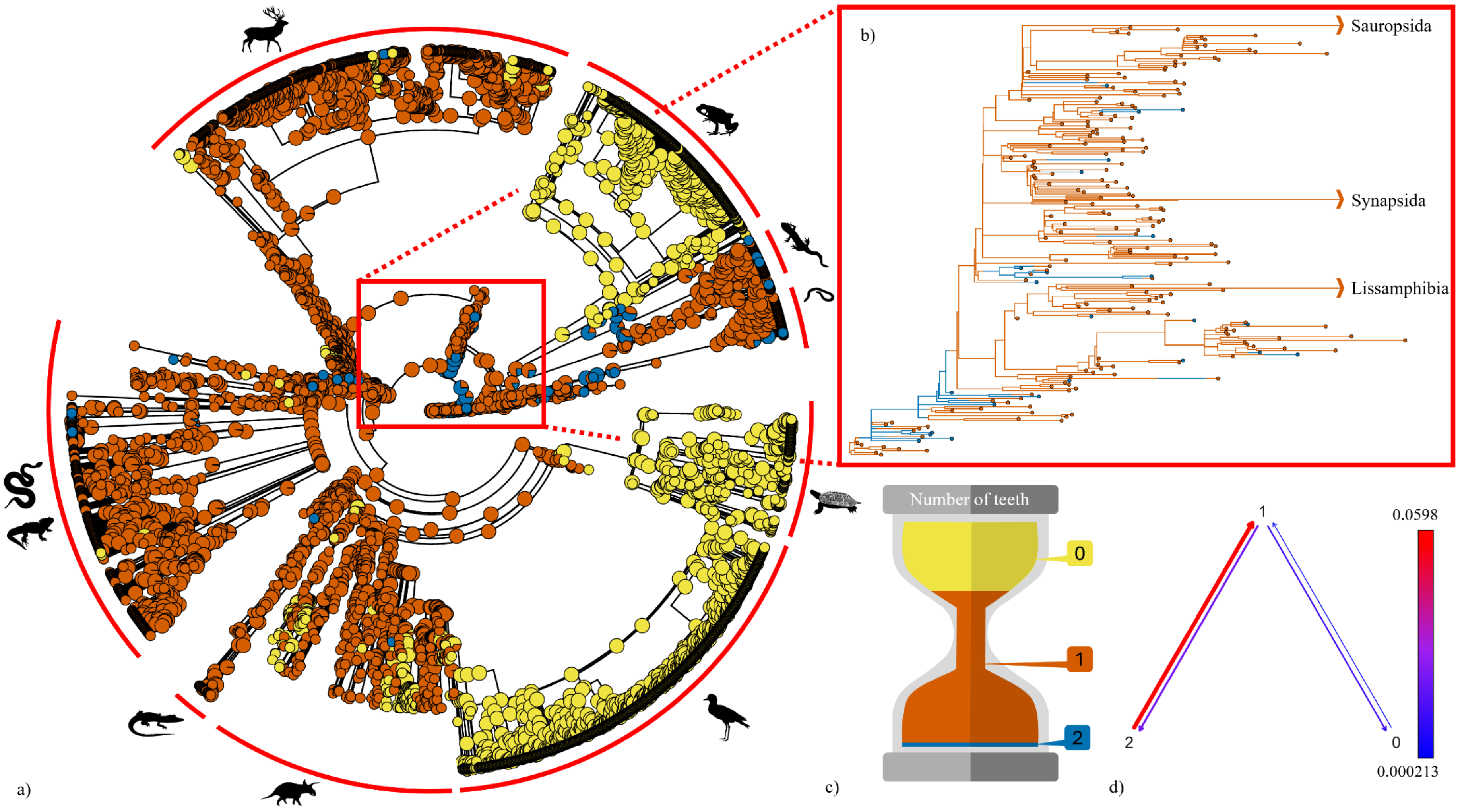

Figure S4: Histogram of number of teeth in the hemimandible.

*Number of specimens = 2611*
*Range of number of teeth in hemimandible = 0:130*

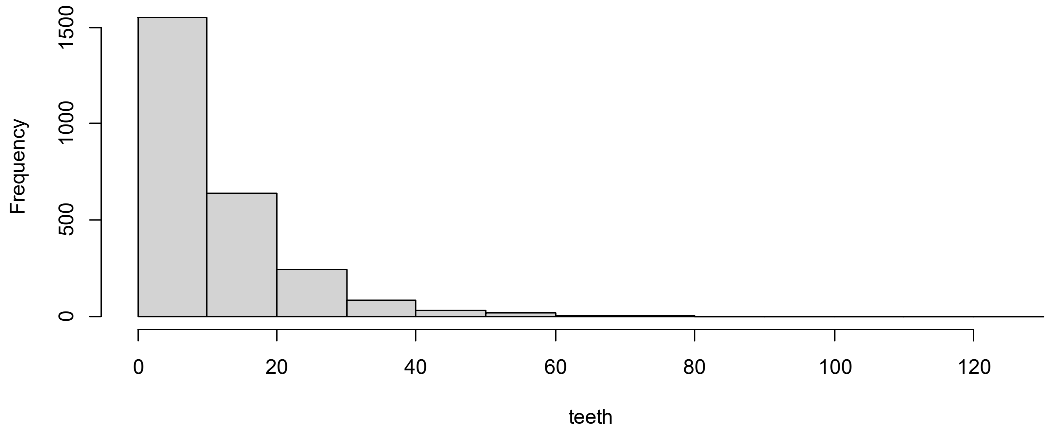

Figure S5: Principal coordinate analysis (PCoA) of all tetrapod jaw characters across the entire dataset with a density background, showing the clustering of points particularly in the extant frogs, birds, and mammals.

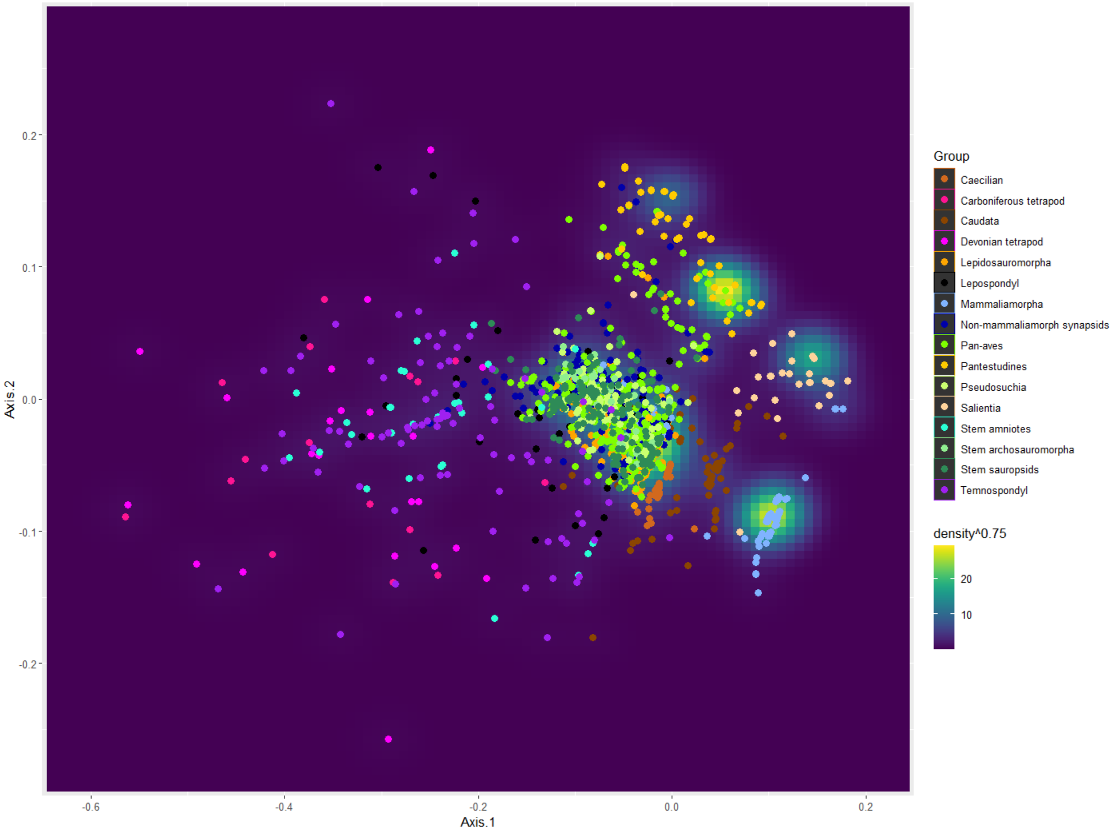

Figure S6: Densities of (a) elements, (b) tooth-bearing elements, and (c) teeth in the hemimandible by time period.

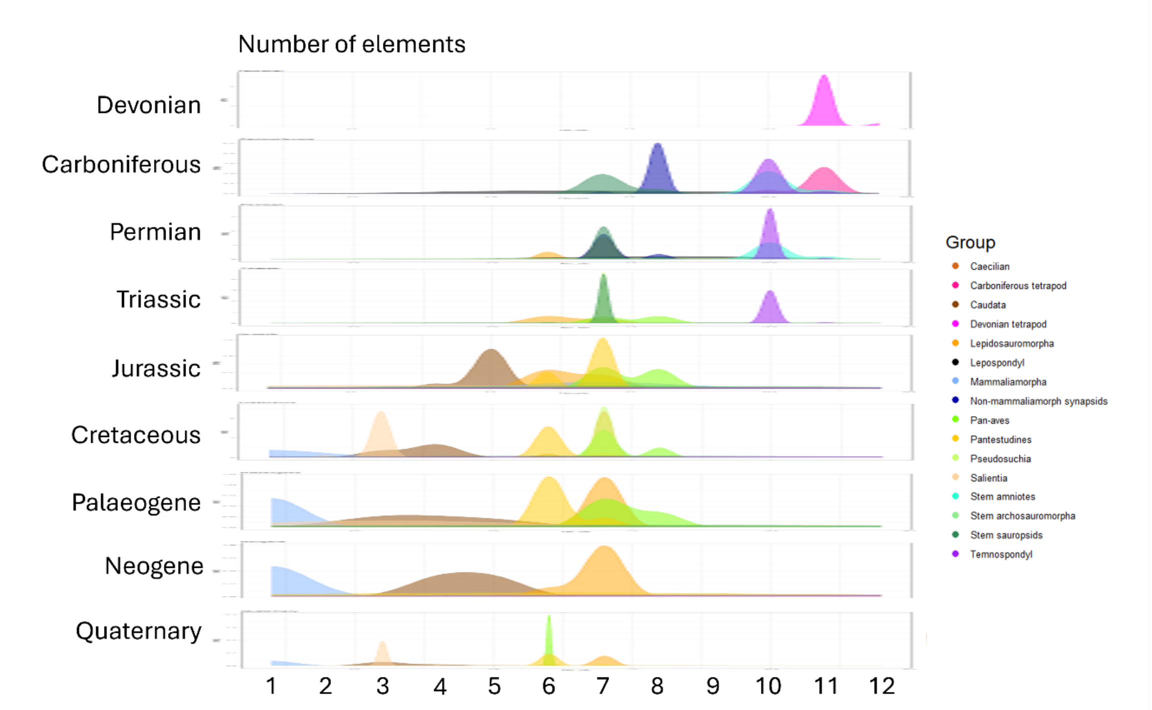

(a)

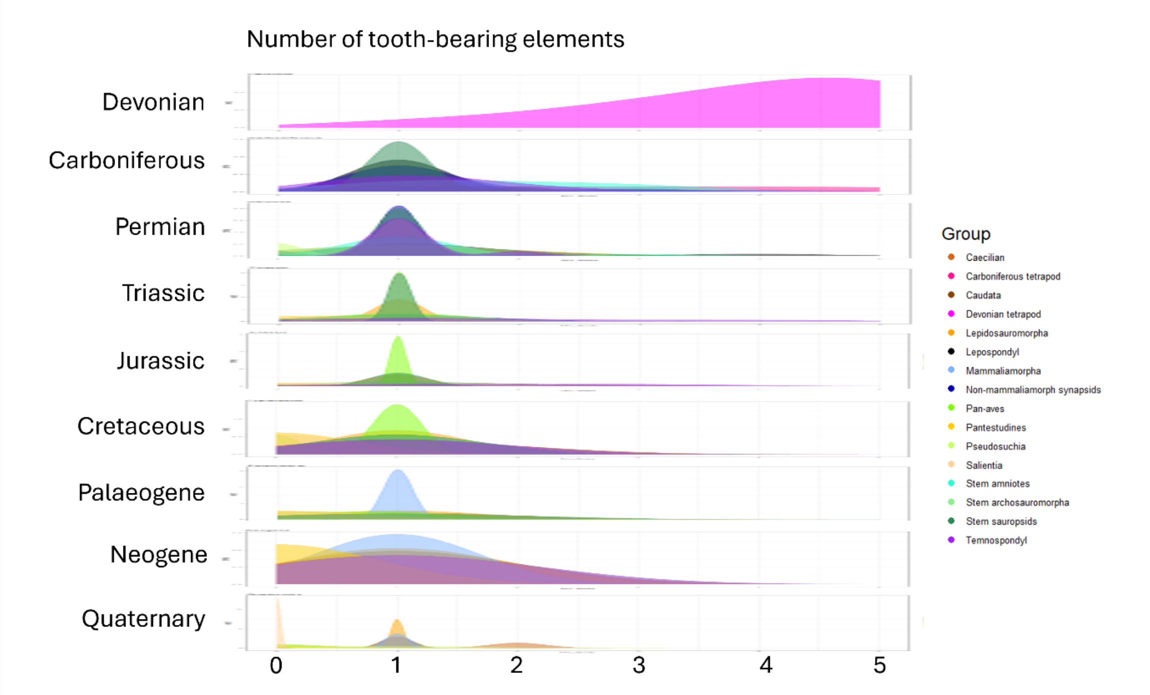

(b)

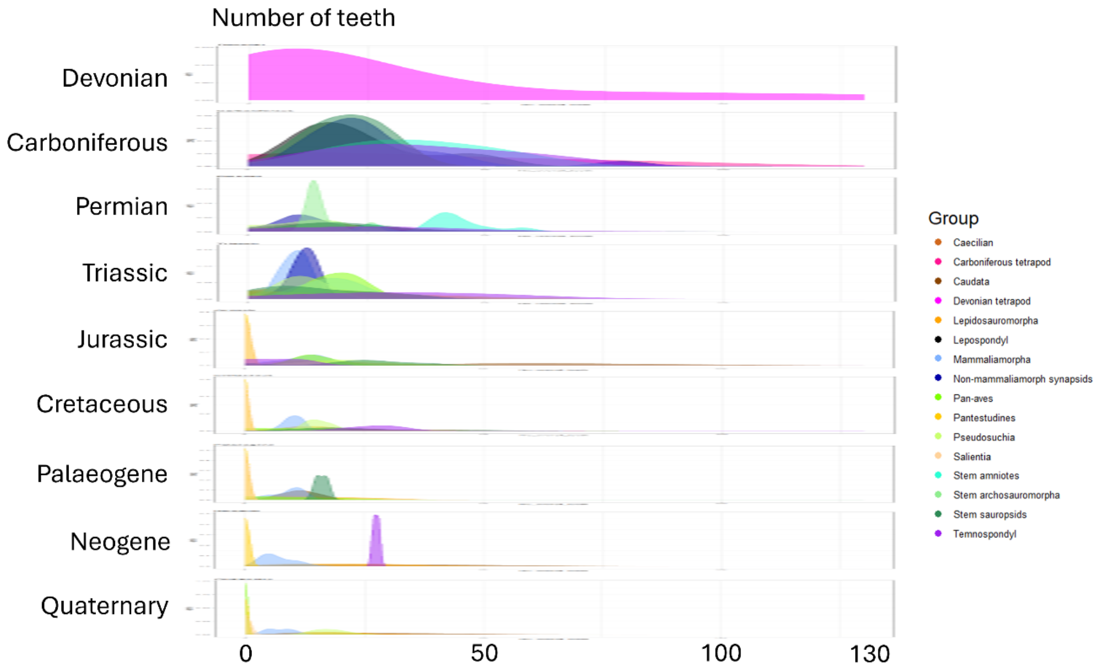

Number of teeth
Devonian
Carboniferous
Permian
Triassic
Jurassic
Cretaceous
Palaeogene
Neogene
Quaternary
0
50
100
130
Group
Caecilian
Carboniferous tetrapod
Caudata
Devonian tetrapod
Lepidosauromorpha
Lepospondyl
Mammaliamorpha
Non-mammaliamorph synapsids
Pan-aves
Pantestudines
Pseudosuchia
Salientia
Stem amniotes
Stem archosauromorpha
Stem sauropsids
Temnospondyl

(c)

Figure S7: Posterior probability distributions of state changes obtained from all-rates-differ (ARD) stochastic mapping of the number of teeth in the hemimandible. HPD = 95% high probability density interval. Three states; 0 = 0 teeth, 1 = 1-50 teeth, 2 = 51+ teeth.

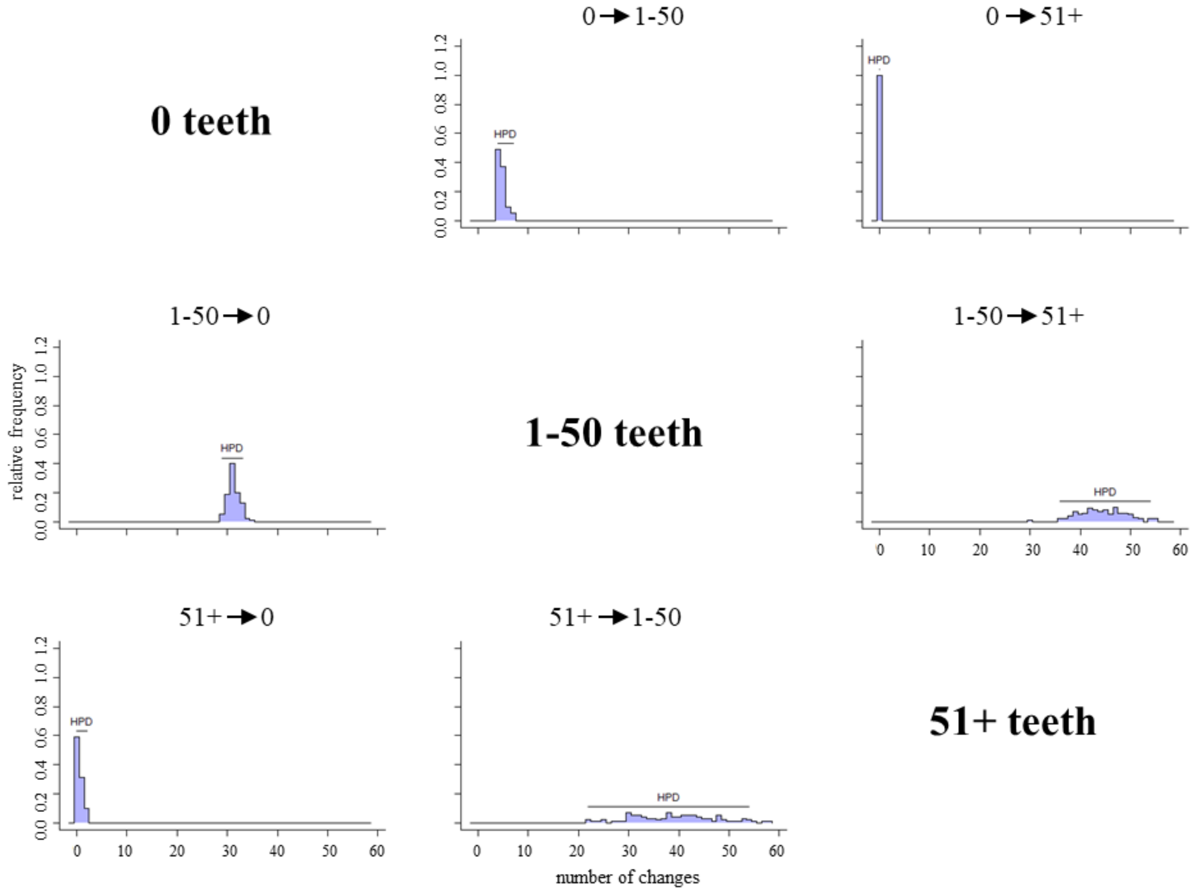

Figure S8: Posterior probability distributions of state changes obtained from all-rates-differ (ARD) stochastic mapping of the number of tooth-bearing elements in the hemimandible. HPD = 95% high probability density interval. Six states; 0-5 tooth-bearing elements (tbels).

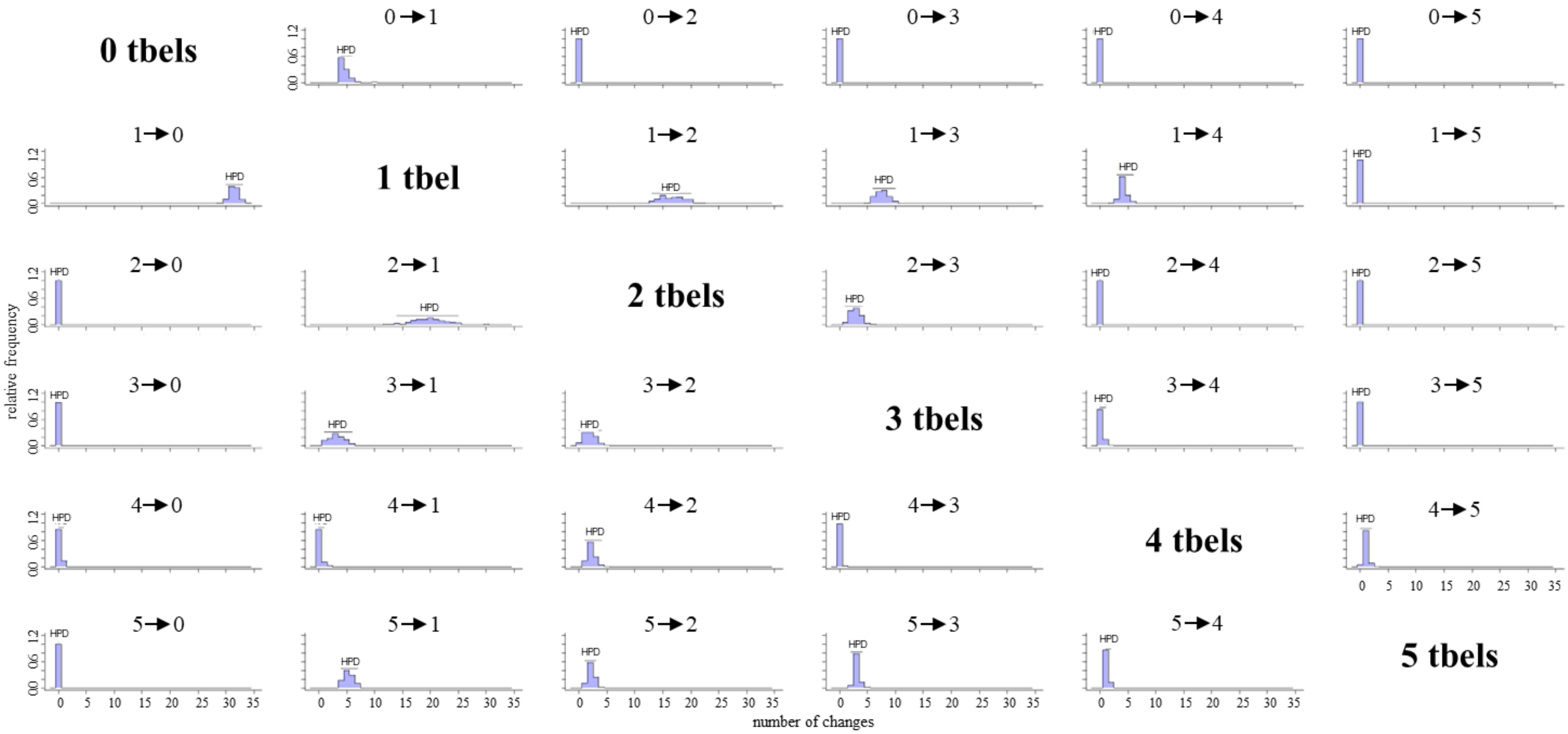

Figure S9: Posterior probability distributions of state changes obtained from symmetric (SYM) stochastic mapping of the number of elements in the hemimandible. HPD = 95% high probability density interval. Twelve states; 1-12 elements (els).

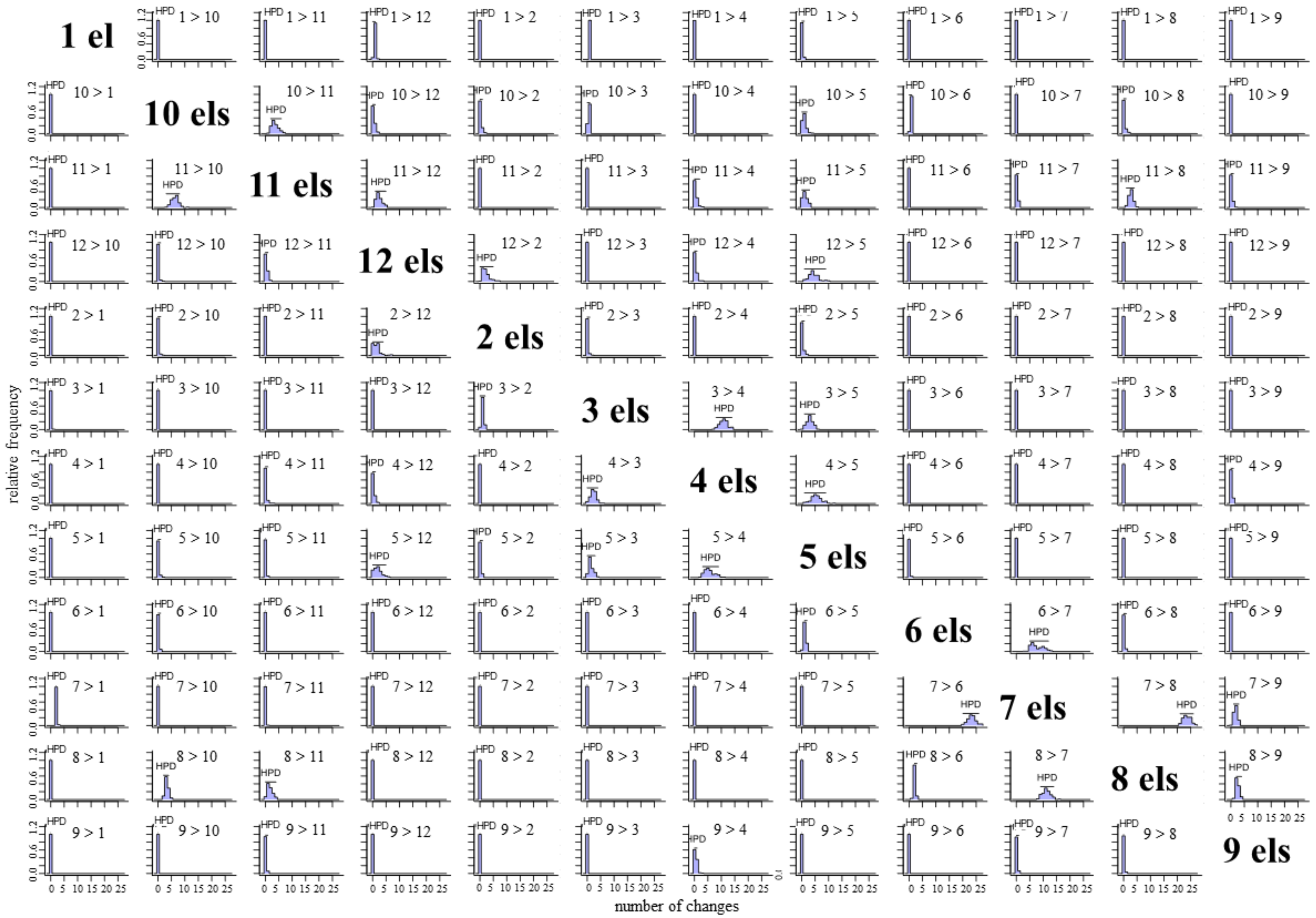

### 2. Supplementary tables

Table S1: AIC scores for fitMk evolutionary modelling. Models with best fit indicated with grey cells.

| | **Equal rates** | **Symmetrical rates** | **Unequal rates** |
|---|---|---|---|
| **Number of teeth** | 928.3707 | 901.2084 | **824.5648** |
| **Number of tooth-bearing elements** | 1221.283 | 1139.197 | **1053.131** |
| **Number of elements** | 1643.163 | **1436.603** | 1494.625 |

Table S2: AICc scores for correlated evolutionary modelling (corHMM). Models with best fit indicated with grey cells.

| | **Teeth vs tooth-bearing elements** | **Teeth vs elements** | **Elements vs tooth-bearing elements** |
|---|---|---|---|
| **Independent ARD** | 1490.194 | 4178.894 | 5371.755 |
| **Correlated ARD** | **1469.065** | 4175.774 | 5347.806 |
| **Independent ERM** | 1478.879 | 4185.41 | 5380.447 |
| **Correlated ERM** | 1488.138 | 4188.712 | 5348.607 |
| **Independent SYM** | 1473.425 | 4171.555 | 5360.694 |
| **Correlated SYM** | 1593.825 | **2998.913** | **3387.561** |

Table S3: PCoA disparity – clade

| Subsets | n | obs | bs.median | 2.50% | 25% | 75% | 97.50% |
|---|---|---|---|---|---|---|---|
| Devonian_eotetrapodiforms | 24 | 0.26 | 0.25 | 0.208 | 0.234 | 0.264 | 0.289 |
| Carboniferous_eotetrapodiforms | 16 | 0.246 | 0.245 | 0.205 | 0.229 | 0.254 | 0.284 |
| Stem_amniotes | 30 | 0.162 | 0.16 | 0.14 | 0.149 | 0.171 | 0.186 |
| Lepospondyls | 40 | 0.151 | 0.149 | 0.131 | 0.14 | 0.154 | 0.166 |
| Temnospondyls | 90 | 0.181 | 0.179 | 0.159 | 0.175 | 0.184 | 0.192 |
| Non_mammaliamorph_synapsids | 200 | 0.087 | 0.086 | 0.081 | 0.084 | 0.088 | 0.092 |
| Mammaliamorpha | 514 | 0.046 | 0.046 | 0.044 | 0.045 | 0.048 | 0.05 |
| Stem_sauropsids | 179 | 0.074 | 0.074 | 0.07 | 0.073 | 0.076 | 0.078 |
| Pantestudines | 154 | 0.085 | 0.083 | 0.048 | 0.08 | 0.085 | 0.105 |
| Lepidosauromorpha | 362 | 0.063 | 0.063 | 0.059 | 0.061 | 0.064 | 0.067 |
| Stem_archosauromorpha | 40 | 0.076 | 0.077 | 0.07 | 0.075 | 0.08 | 0.085 |
| Pseudosuchia | 127 | 0.085 | 0.085 | 0.081 | 0.083 | 0.086 | 0.087 |
| Pan_aves | 874 | 0.084 | 0.084 | 0.075 | 0.082 | 0.087 | 0.088 |
| Salientia | 263 | 0.016 | 0.017 | 0.012 | 0.015 | 0.018 | 0.021 |
| Caudata | 85 | 0.116 | 0.116 | 0.095 | 0.109 | 0.12 | 0.128 |
| Gymnophiona | 49 | 0.044 | 0.043 | 0.035 | 0.039 | 0.048 | 0.061 |

Table S4: PCoA disparity – time period

| subsets | n | obs | bs.median | 2.50% | 25% | 75% | 97.50% |
|---|---|---|---|---|---|---|---|
| Devonian | 24 | 0.26 | 0.251 | 0.197 | 0.231 | 0.262 | 0.281 |
| Carboniferous | 96 | 0.163 | 0.163 | 0.141 | 0.157 | 0.167 | 0.179 |
| Permian | 307 | 0.105 | 0.104 | 0.098 | 0.102 | 0.107 | 0.112 |
| Triassic | 306 | 0.093 | 0.093 | 0.089 | 0.091 | 0.094 | 0.096 |
| Jurassic | 174 | 0.09 | 0.09 | 0.081 | 0.087 | 0.092 | 0.096 |
| Cretaceous | 467 | 0.101 | 0.101 | 0.095 | 0.099 | 0.103 | 0.107 |
| Palaeogene | 149 | 0.105 | 0.107 | 0.092 | 0.101 | 0.112 | 0.124 |
| Neogene | 58 | 0.11 | 0.107 | 0.074 | 0.1 | 0.116 | 0.128 |
| Quaternary | 1466 | 0.109 | 0.11 | 0.107 | 0.108 | 0.111 | 0.115 |

Table S5: PCoA disparity – time period significance testing

Permutation test for adonis under reduced model
Permutation: free
Number of permutations: 999

| | Df | SumOfSqs | R2 | F | Pr(>F) |
|---|---|---|---|---|---|
| Model | 8 | -0.0159 | -0.0015 | -0.5682 | 1 |
| Residual | 3038 | 10.6511 | 1.0015 | | |
| Total | 3046 | 10.6351 | 1 | | |

Table S6: Summary of all-rates-differ (ARD) stochastic character mapping of number of teeth in the hemimandible (n = 100 iterations).

States: zero teeth (0), 1-50 teeth (1), and 51+ teeth (2)

Trees have 120 changes between states on average.

Distribution of changes from stochastic mapping:
Cumulative increases: min. 34, max. 62
Cumulative decreases: min. 51, max. 95

Mean total time spent in each state

| | 0 | 1 | 2 | total |
|---|---|---|---|---|
| Time | 2.149663e+04 | 4.720065e+04 | 1.405314e+03 | 70102.59 |
| Proportion | 3.066453e-01 | 6.733081e-01 | 2.004653e-02 | 1.00 |

Table S7: Summary of all-rates-differ (ARD) stochastic character mapping of number of tooth-bearing elements in the hemimandible (n = 100 iterations).

States: zero tooth-bearing elements (0), one tooth-bearing element (1), two tooth-bearing elements (2), three tooth-bearing elements (3), four tooth-bearing elements (4), five tooth-bearing elements (5).

Trees have 108.13 changes between states on average.

Distribution of changes from stochastic mapping:
Cumulative increases: min. 25, max. 56
Cumulative decreases: min. 48, max. 107

Mean total time spent in each state

| | 0 | 1 | 2 | 3 | 4 | 5 | total |
|---|---|---|---|---|---|---|---|
| Time | 2.149700e+04 | 4.557247e+04 | 2.24404e+03 | 2.252557e+02 | 2.206148e+02 | 3.432082e+02 | 70102.59 |
| Proportion | 3.066506e-01 | 6.500826e-01 | 3.20108e-02 | 3.213229e-03 | 3.147028e-03 | 4.895799e-03 | 1.00 |

Table S8: Summary of symmetric (SYM) stochastic character mapping of number of elements in the hemimandible (n = 100 iterations).

States: one element (1), two elements (2), three elements (3), four elements (4), five elements (5), six elements (6), seven elements (7), eight elements (8), nine elements (9), ten elements (10), eleven elements (11), twelve elements (12).

Trees have 144.84 changes between states on average.

Distribution of changes from stochastic mapping:
Cumulative increases: min. 36, max. 141
Cumulative decreases: min., 38 max. 138

Mean total time spent in each state

| | 1 | 2 | 3 | 4 | 5 | 6 |
|---|---|---|---|---|---|---|
| Time | 8497.1606149 | 32.797635433 | 9278.6587420 | 746.39244323 | 721.25859211 | 1.300534e+04 |
| Proportion | 0.1212104 | 0.000467852 | 0.1323583 | 0.01064714 | 0.01028861 | 1.855186e-01 |

| | 7 | 8 | 9 | 10 | 11 | 12 | total |
|---|---|---|---|---|---|---|---|
| Time | 2.973067e+04 | 4.832802e+03 | 1.247923e+02 | 2.255752e+03 | 798.10541879 | 78.869996990 | 70102.59 |
| Proportion | 4.241022e-01 | 6.893898e-02 | 1.780139e-03 | 3.217787e-02 | 0.01138482 | 0.001125065 | 1.00 |

### 3. Composite tree references

Abello, M.A. & Candela, A.M. (2020) Paleobiology of *Argyrolagus* (Marsupialia, Argyrolagidae): an astonishing case of bipedalism among South American mammals. *Journal of Mammalian Evolution*, **27**, 419–444.

Agnolin, F. (2012) A new Calyptocephalellidae (Anura, Neobatrachia) from the Upper Cretaceous of Patagonia, Argentina, with comments on its systematic position. *Studia geologica salmanticensia*, **48**, 129–178.

Agnolin, F.L., Gonzalez Riga, B.J., Aranciaga Rolando, A.M., Rozadilla, S., Motta, M.J., Chimento, N.R., *et al.* (2023) A new giant titanosaur (Dinosauria, Sauropoda) from the Upper Cretaceous of Northwestern Patagonia, Argentina. *Cretaceous Research*, **146**, 105487.

Agnolin, F.L., Motta, M.J., Brissón Egli, F., Lo Coco, G. & Novas, F.E. (2019) Paravian Phylogeny and the Dinosaur-Bird Transition: An Overview. *Frontiers in Earth Science*, **6**.

Alifanov, V.R. (2012) Lizards of the family Arretosauridae Gilmore, 1943 (Iguanomorpha, Iguania) from the Paleogene of Mongolia. *Paleontological Journal*, **46**, 412–420.

Alifanov, V.R. (2016) Lizards of the family Hodzhakuliidae (Scincomorpha) from the lower Cretaceous of Mongolia. *Paleontological Journal*, **50**, 504–513.

Alifanov, V.R. (2018) Lizards of the Family Temujiniidae (Iguanomorpha): Finds from the Aptian–Albian of Mongolia, Classification and Geographical Origin. *Paleontological Journal*, **52**, 653–663.

Alifanov, V.R. (2019) Lizards of the Families Eoxantidae, Ardeosauridae, Globauridae, and Paramacellodidae (Scincomorpha) from the Aptian–Albian of Mongolia. *Paleontological Journal*, **53**, 74–88.

Anderson, J.S. (2003) A new aïstopod (Tetrapoda: Lepospondyli) from Mazon Creek, Illinois. *Journal of Vertebrate Paleontology*, **23**, 79–88.

Andres, B. (2021) Phylogenetic systematics of *Quetzalcoatlus* Lawson 1975 (Pterodactyloidea:Azhdarchoidea). *Journal of Vertebrate Paleontology*, **41**, 203–217.

Andres, B., Clark, J. & Xu, X. (2014) The Earliest Pterodactyloid and the Origin of the Group. *Current Biology*, **24**, 1011–1016.

Andrews, S.M. & Carroll, R.L. (1991) The Order Adelospondyli: Carboniferous lepospondyl amphibians. *Earth and Environmental Science Transactions of The Royal Society of Edinburgh*, **82**, 239–275.

Andrzejewski, K.A., Winkler, D.A. & Jacobs, L.L. (2019) A new basal ornithopod (Dinosauria: Ornithischia) from the Early Cretaceous of Texas. *PLOS ONE*, **14**, e0207935.

Angielczyk, K.D., Liu, J. & Yang, W. (2021) A redescription of *Kunpania scopulusa* , a bidentalian dicynodont (Therapsida, Anomodontia) from the ?Guadalupian of northwestern China. *Journal of Vertebrate Paleontology*, **41**, e1922428.

Angielczyk, K.D. & Ruta, M. (2012) The Roots of Amphibian Morphospace: A Geometric Morphometric Analysis of Paleozoic Temnospondyls. *Fieldiana Life and Earth Sciences*, **2012**, 40–58.

Apesteguía, S., Gómez, R.O. & Rougier, G.W. (2012) A basal sphenodontian (Lepidosauria) from the Jurassic of Patagonia: new insights on the phylogeny and biogeography of Gondwanan rhynchocephalians. *Zoological Journal of the Linnean Society*, **166**, 342–360.

Arbour, V.M. & Currie, P.J. (2016) Systematics, phylogeny and palaeobiogeography of the ankylosaurid dinosaurs. *Journal of Systematic Palaeontology*, **14**, 385–444.

Atayman, S., Rubidge, B.S. & Abdala, F. (2009) Taxonomic re-evaluation of tapinocephalid dinocephalians. In *15th Biennial Meeting of the Palaeontological Society of Southern Africa*. Presented at the 15th Biennial Meeting of the Palaeontological Society of Southern Africa, Palaeontologia Africana, pp. 88–90.

Averianov, A. & Lopatin, A. (2021) A New Theropod Dinosaur (Theropoda, Dromaeosauridae) from the Late Cretaceous of Tajikistan. *Doklady Earth Sciences*, **499**, 570–574.

Averianov, A. & Sues, H.-D. (2016) Troodontidae (Dinosauria: Theropoda) from the Upper Cretaceous of Uzbekistan. *Cretaceous Research*, **59**, 98–110.

Averianov, A.O., Lopatin, A.V. & Leshchinskiy, S.V. (2023) New interpretation of dentition in Early Cretaceous docodontan *Sibirotherium* based on micro-computed tomography. *Journal of Mammalian Evolution*.

Ballell, A., Moon, B.C., Porro, L.B., Benton, M.J. & Rayfield, E.J. (2019) Convergence and functional evolution of longirostry in crocodylomorphs. *Palaeontology*, **62**, 867–887.

Bandeira, K.L.N., Simbras, F.M., Machado, E.B., Campos, D. de A., Oliveira, G.R. & Kellner, A.W.A. (2016) A New Giant Titanosauria (Dinosauria: Sauropoda) from the Late Cretaceous Bauru Group, Brazil. *PLOS ONE*, **11**, e0163373.

Baron, M.G., Norman, D.B. & Barrett, P.M. (2017) A new hypothesis of dinosaur relationships and early dinosaur evolution. *Nature*, **543**, 501–506.

Barrett, P.Z. (2021) The largest hoplophonine and a complex new hypothesis of nimravid evolution. *Scientific Reports*, **11**, 21078.

Barrett, P.Z., Hopkins, S.S.B. & Price, S.A. (2021) How many sabertooths? Reevaluating the number of carnivoran sabretooth lineages with total-evidence Bayesian techniques and a novel origin of the Miocene Nimravidae. *Journal of Vertebrate Paleontology*, **41**, e1923523.

Barrientos-Lara, J.I. & Alvarado-Ortega, J. (2020) *Acuetzpalin carranzai* gen et sp. nov. A new ophthalmosauridae (Ichthyosauria) from the Upper Jurassic of Durango, North Mexico. *Journal of South American Earth Sciences*, **98**, 102456.

Barrientos-Lara, J.I. & Alvarado-Ortega, J. (2021) A new ophthalmosaurid (Ichthyosauria) from the Upper Kimmeridgian deposits of the La Casita Formation, near Gómez Farías, Coahuila, northern Mexico. *Journal of South American Earth Sciences*, **111**, 103499.

Beccari, V., Mateus, O., Wings, O., Milàn, J. & Clemmensen, L.B. (2021) *Issi saaneq* gen. et sp. nov.—A New Sauropodomorph Dinosaur from the Late Triassic (Norian) of Jameson Land, Central East Greenland. *Diversity*, **13**, 561.

Beck, R.M.D. (2023) Diversity and Phylogeny of Marsupials and Their Stem Relatives (Metatheria). In *American and Australasian Marsupials: An Evolutionary, Biogeographical, and Ecological Approach* (ed. by Cáceres, N.C. & Dickman, C.R.). Springer International Publishing, Cham, pp. 1–66.

Beck, R.M.D., Voss, R.S. & Jansa, S.A. (2022) Craniodental Morphology and Phylogeny of Marsupials. *Bulletin of the American Museum of Natural History*, **457**, 1–352.

Benson, R.B.J., Evans, M. & Druckenmiller, P.S. (2012) High Diversity, Low Disparity and Small Body Size in Plesiosaurs (Reptilia, Sauropterygia) from the Triassic–Jurassic Boundary. *PLOS ONE*, **7**, e31838.

Berman, D.S., Maddin, H.C., Henrici, A.C., Sumida, S.S., Scott, D. & Reisz, R.R. (2020) New Primitive Caseid (Synapsida, Caseasauria) from the Early Permian of Germany. *Annals of Carnegie Museum*, **86**, 43–75.

Beznosov, P.A., Clack, J.A., Lukševičs, E., Ruta, M. & Ahlberg, P.E. (2019) Morphology of the earliest reconstructable tetrapod *Parmastega aelidae*. *Nature*, **574**, 527–531.

Bishop, P.J. & Pierce, S.E. (2023) The fossil record of appendicular muscle evolution in Synapsida on the line to mammals: Part I—Forelimb. *The Anatomical Record*, **n/a**.

Blanco, A. (2021) Importance of the postcranial skeleton in eusuchian phylogeny: Reassessing the systematics of allodaposuchid crocodylians. *PLOS ONE*, **16**, e0251900.

Blieck, A., Clément, G. & Streel, M. (2010) The biostratigraphical distribution of earliest tetrapods (Late Devonian): a revised version with comments on biodiversification. *Geological Society, London, Special Publications*, **339**, 129–138.

Boessenecker, R.W. & Fordyce, R.E. (2017) A new eomysticetid from the Oligocene Kokoamu Greensand of New Zealand and a review of the Eomysticetidae (Mammalia, Cetacea). *Journal of Systematic Palaeontology*, **15**, 429–469.

Bolet, A., Stubbs, T.L., Herrera-Flores, J.A. & Benton, M.J. (2022) The Jurassic rise of squamates as supported by lepidosaur disparity and evolutionary rates. *eLife*, **11**, e66511.

Bonaparte, J.F. (2013) Evolution of the Brasilodontidae (Cynodontia-Eucynodontia). *Historical Biology*, **25**, 643–653.

Boscaini, A., Pujos, F. & Gaudin, T.J. (2019) A reappraisal of the phylogeny of Mylodontidae (Mammalia, Xenarthra) and the divergence of mylodontine and lestodontine sloths. *Zoologica Scripta*, **48**, 691–710.

Bourque, J.R., Howard Hutchison, J., Holroyd, P.A. & Bloch, J.I. (2015) A new dermatemydid (Testudines, Kinosternoidea) from the Paleocene-Eocene Thermal Maximum, Willwood Formation, southeastern Bighorn Basin, Wyoming. *Journal of Vertebrate Paleontology*, **35**, e905481.

Bourque, J.R. & Schubert, B.W. (2015) Fossil musk turtles (Kinosternidae, Sternotherus) from the late Miocene–early Pliocene (Hemphillian) of Tennessee and Florida. *Journal of Vertebrate Paleontology*, **35**, e885441.

Bredehoeft, K.E. & Schubert, B.W. (2015) A Re-Evaluation of the Pleistocene Hellbender, *Cryptobranchus guildayi*. *Journal of Herpetology*, **49**, 157–160.

Brinkman, D.B., Densmore, M., Rabi, M., Ryan, M.J. & Evans, D.C. (2015) Marine turtles from the Late Cretaceous of Alberta, Canada. *Canadian Journal of Earth Sciences*, **52**, 581–589.

Brinkman, D.B. & Eberth, D.A. (1984) A new arceoscelid reptile, *Zarcasaurus tanyderus*, from the Gutler Formation (Lower Permian) of north-central New Mexico. *New Mexico Geology*, 34–39.

Britt, B.B., Dalla Vecchia, F.M., Chure, D.J., Engelmann, G.F., Whiting, M.F. & Scheetz, R.D. (2018) *Caelestiventus hanseni* gen. et sp. nov. extends the desert-dwelling pterosaur record back 65 million years. *Nature Ecology & Evolution*, **2**, 1386–1392.

Brum, A.S., Pêgas, R.V., Bandeira, K.L.N., Souza, L.G., Campos, D.A. & Kellner, A.W.A. (2021) A new unenlagiine (Theropoda, Dromaeosauridae) from the Upper Cretaceous of Brazil. *Papers in Palaeontology*, **7**, 2075–2099.

Brusatte, S.L., Benson, R.B.J. & Xu, X. (2012) A Reassessment of *Kelmayisaurus petrolicus*, a Large Theropod Dinosaur from the Early Cretaceous of China. *Acta Palaeontologica Polonica*, **57**, 65–72.

Brust, A.C.B., Desojo, J.B., Schultz, C.L., Paes-Neto, V.D. & Da-Rosa, Á.A.S. (2018) Osteology of the first skull of *Aetosauroides scagliai* Casamiquela 1960 (Archosauria: Aetosauria) from the Upper Triassic of southern Brazil (*Hyperodapedon* Assemblage Zone) and its phylogenetic importance. *PLOS ONE*, **13**, e0201450.

Buffa, V., Jalil, N.-E. & Steyer, J.-S. (2019) Redescription of *Arganasaurus* (*Metoposaurus*) *azerouali* (Dutuit) comb. nov. from the Upper Triassic of the Argana Basin (Morocco), and the first phylogenetic analysis of the Metoposauridae (Amphibia, Temnospondyli). *Papers in Palaeontology*, **5**, 699–717.

Buscalioni, Á.D. (2017) The Gobiosuchidae in the early evolution of Crocodyliformes. *Journal of Vertebrate Paleontology*, **37**, e1324459.

Butler, R.J., Ezcurra, M.D., Montefeltro, F.C., Samathi, A. & Sobral, G. (2015) A new species of basal rhynchosaur (Diapsida: Archosauromorpha) from the early Middle Triassic of South Africa, and the early evolution of Rhynchosauria: New basal rhynchosaur from South Africa. *Zoological Journal of the Linnean Society*, **174**, 571–588.

Butler, R.J., Fernandez, V., Nesbitt, S.J., Leite, J.V. & Gower, D.J. (2022) A new pseudosuchian archosaur, *Mambawakale ruhuhu* gen. et sp. nov., from the Middle Triassic Manda Beds of Tanzania. *Royal Society Open Science*, **9**, 211622.

Cadena, E.-A., Scheyer, T.M., Carrillo-Briceño, J.D., Sánchez, R., Aguilera-Socorro, O.A., Vanegas, A., *et al.* (2020) The anatomy, paleobiology, and evolutionary relationships of the largest extinct side-necked turtle. *Science Advances*, **6**, eaay4593.

Calvo, J.O. & Riga, B.G. (2018) *Baalsaurus mansillai* gen. et sp. nov. a new titanosaurian sauropod (Late Cretaceous) from Neuquén, Patagonia, Argentina. *Anais da Academia Brasileira de Ciências*, **91**.

Canudo, J., Carballido, J., Salgado, L. & Garrido, A. (2018) A new rebbachisaurid sauropod from the Aptian–Albian, Lower Cretaceous Rayoso Formation, Neuquén, Argentina. *Acta Palaeontologica Polonica*, **63**, 679–691.

Cardillo, M., Bininda-Emonds, O.R.P., Boakes, E. & Purvis, A. (2004) A species-level phylogenetic supertree of marsupials. *Journal of Zoology*, **264**, 11–31.

Carlson, K.J. (1999) *Crossotelos*, an Early Permian nectridian amphibian. *Journal of Vertebrate Paleontology*, **19**, 623–631.

Carr, T.D., Varricchio, D.J., Sedlmayr, J.C., Roberts, E.M. & Moore, J.R. (2017) A new tyrannosaur with evidence for anagenesis and crocodile-like facial sensory system. *Scientific Reports*, **7**, 44942.

Carr, T.D. & Williamson, T.E. (2010) *Bistahieversor sealeyi* , gen. et sp. nov., a new tyrannosauroid from New Mexico and the origin of deep snouts in Tyrannosauroidea. *Journal of Vertebrate Paleontology*, **30**, 1–16.

Carrano, M.T., Benson, R.B.J. & Sampson, S.D. (2012) The phylogeny of Tetanurae (Dinosauria: Theropoda). *Journal of Systematic Palaeontology*, **10**, 211–300.

Carvalho, I.S., Agnolin, F., Aranciaga Rolando, M.A., Novas, F.E., Xavier-Neto, J., Freitas, F.I., *et al.* (2019) A new genus of pipimorph frog (Anura) from the Early Cretaceous Crato Formation (Aptian) and the evolution of South American tongueless frogs. *Journal of South American Earth Sciences*, **92**, 222–233.

Cerda, I., Zurriaguz, V.L., Carballido, J.L., González, R. & Salgado, L. (2021) Osteology, paleohistology and phylogenetic relationships of Pellegrinisaurus powelli (Dinosauria:

Sauropoda) from the Upper Cretaceous of Argentinean Patagonia. *Cretaceous Research*, **128**, 104957.

Čerňanský, A. (2019) The first potential fossil record of a dibamid reptile (Squamata: Dibamidae): a new taxon from the early Oligocene of Central Mongolia. *Zoological Journal of the Linnean Society*, **187**, 782–799.

Čerňanský, A., Augé, M. louis & Rage, J. (2015) A complete mandible of a new amphisbaenian reptile (Squamata, Amphisbaenia) from the late middle eocene (Bartonian, Mp 16) of France. *Journal of Vertebrate Paleontology*, **35**, e902379.

Čerňanský, A., Bolet, A., Müller, J., Rage, J.-C. & Herrel, A. (2017) A new exceptionally preserved specimen of *Dracaenosaurus* (Squamata, Lacertidae) from the Oligocene of France as revealed by micro-computed tomography. *Journal of Vertebrate Paleontology*, **37**, e1384738.

Chambi-Trowell, S.A.V., Martinelli, A.G., Whiteside, D.I., Vivar, P.R.R. de, Soares, M.B., Schultz, C.L., *et al.* (2021) The diversity of Triassic South American sphenodontians: a new basal form, clevosaurs, and a revision of rhynchocephalian phylogeny. *Journal of Systematic Palaeontology*, **19**, 787–820.

Chapelle, K.E.J., Barrett, P.M., Botha, J. & Choiniere, J.N. (2019) *Ngwevu intloko*: a new early sauropodomorph dinosaur from the Lower Jurassic Elliot Formation of South Africa and comments on cranial ontogeny in *Massospondylus carinatus*. *PeerJ*, **7**, e7240.

Chen, C., Xiaoyou, H., Wei, L., Lingyun, Y., Haigang, C. & Xinping, Z. (2021) Complete mitochondrial genome and the phylogenetic position of the Burmese narrow-headed softshell turtle *Chitra vandijki* (Testudines: Trionychidae). *Mitochondrial DNA Part B*, **6**, 1216–1218.

Chen, D., Alavi, Y., Brazeau, M., Blom, H., Millward, D. & Ahlberg, P. (2017) A partial lower jaw of a tetrapod from "Romer's Gap." *Earth and Environmental Science Transactions of the Royal Society of Edinburgh*, **108**, 55–65.

Chen, J., Bever, G.S., Yi, H.-Y. & Norell, M.A. (2016) A burrowing frog from the late Paleocene of Mongolia uncovers a deep history of spadefoot toads (Pelobatoidea) in East Asia. *Scientific Reports*, **6**, 19209.

Chimento, N.R., Agnolín, F.L., Manabe, M., Tsuihiji, T., Rich, T.H., Vickers-Rich, P., *et al.* (2023) First monotreme from the Late Cretaceous of South America. *Communications Biology*, **6**, 1–6.

Choiniere, J.N., Forster, C.A. & Klerk, W.J. de. (2012) New information on *Nqwebasaurus thwazi*, a coelurosaurian theropod from the Early Cretaceous Kirkwood Formation in South Africa. *Journal of African Earth Sciences*, **71–72**, 1–17.

Chokchaloemwong, D., Hattori, S., Cuesta, E., Jintasakul, P., Shibata, M. & Azuma, Y. (2019) A new carcharodontosaurian theropod (Dinosauria: Saurischia) from the Lower Cretaceous of Thailand. *PLOS ONE*, **14**, e0222489.

Chongxi, Y. (2008) A New Genus and Species of Sapeornithidae from Lower Cretaceous in Western Liaoning, China. *Acta Geologica Sinica - English Edition*, **82**, 48–55.

Cisneros, J.C. (2008) Phylogenetic relationships of procolophonid parareptiles with remarks on their geological record. *Journal of Systematic Palaeontology*, **6**, 345–366.

Cisneros, J.C., Angielczyk, K., Kammerer, C.F., Smith, R.M.H., Fröbisch, J., Marsicano, C.A., *et al.* (2020a) Captorhinid reptiles from the lower Permian Pedra de Fogo Formation, Piauí, Brazil: the earliest herbivorous tetrapods in Gondwana. *PeerJ*, **8**, e8719.

Cisneros, J.C., Kammerer, C.F., Angielczyk, K.D., Fröbisch, J., Marsicano, C., Smith, R.M.H., *et al.* (2020b) A new reptile from the lower Permian of Brazil (*Karutia fortunata* gen. et sp. nov.) and the interrelationships of Parareptilia. *Journal of Systematic Palaeontology*, **18**, 1939–1959.

Clack, J.A., Bennett, C.E., Carpenter, D.K., Davies, S.J., Fraser, N.C., Kearsey, T.I., *et al.* (2017) Phylogenetic and environmental context of a Tournaisian tetrapod fauna. *Nature Ecology & Evolution*, **1**, 1–11.

Clark, J.M., Sues, H.-D. & Berman, D.S. (2001) A new specimen of *Hesperosuchus agilis* from the Upper Triassic of New Mexico and the interrelationships of basal crocodylomorph archosaurs. *Journal of Vertebrate Paleontology*, **20**, 683–704.

Clément, G. & Lebedev, O. (2014) Revision of the early tetrapod *Obruchevichthys* Vorobyeva, 1977 from the Frasnian (Upper Devonian) of the North-western East European Platform. *Paleontological Journal*, **48**, 1082–1091.

Conrad, J.L. (2015) A New Eocene Casquehead Lizard (Reptilia, Corytophanidae) from North America. *PLOS ONE*, **10**, e0127900.

Conrad, J.L. (2018) A new lizard (Squamata) was the last meal of *Compsognathus* (Theropoda: Dinosauria) and is a holotype in a holotype. *Zoological Journal of the Linnean Society*, **183**, 584–634.

Conrad, J.L., Ast, J.C., Montanari, S. & Norell, M.A. (2011) A combined evidence phylogenetic analysis of Anguimorpha (Reptilia: Squamata). *Cladistics*, **27**, 230–277.

Conrad, J.L., Rieppel, O. & Grande, L. (2007) A Green River (Eocene) polychrotid (Squamata: Reptilia) and a re-examination of iguanian systematics. *Journal of Paleontology*, **81**, 1365–1373.

Coria, R.A. & Currie, P.J. (2016) A New Megaraptoran Dinosaur (Dinosauria, Theropoda, Megaraptoridae) from the Late Cretaceous of Patagonia. *PLOS ONE*, **11**, e0157973.

Coria, R.A., Currie, P.J., Ortega, F. & Baiano, M.A. (2020) An Early Cretaceous, medium-sized carcharodontosaurid theropod (Dinosauria, Saurischia) from the Mulichinco Formation (upper Valanginian), Neuquén Province, Patagonia, Argentina. *Cretaceous Research*, **111**, 104319.

Cozzuol, M.A., Mothé, D. & Avilla, L.S. (2012) A critical appraisal of the phylogenetic proposals for the South American Gomphotheriidae (Proboscidea: Mammalia). *Quaternary International*, Mammoths and Their Relatives 1: Biotopes, Evolution and Human Impact V International Conference, Le Puy-en-Velay, 2010, **255**, 36–41.

Damiani, R.J. (2001) A systematic revision and phylogenetic analysis of Triassic mastodonsauroids (Temnospondyli: Stereospondyli). *Zoological Journal of the Linnean Society*, **133**, 379–482.

Datta, D., Ray, S. & Bandyopadhyay, S. (2021) Cranial morphology of a new phytosaur (Diapsida, Archosauria) from the Upper Triassic of India: implications for phytosaur phylogeny and biostratigraphy. *Papers in Palaeontology*, **7**, 675–708.

Daza, J.D., Stanley, E.L., Wagner, P., Bauer, A.M. & Grimaldi, D.A. (2016) Mid-Cretaceous amber fossils illuminate the past diversity of tropical lizards. *Science Advances*, **2**, e1501080.

Delsuc, F., Gibb, G.C., Kuch, M., Billet, G., Hautier, L., Southon, J., *et al.* (2016) The phylogenetic affinities of the extinct glyptodonts. *Current Biology*, **26**, R155–R156.

DeMar, D.G., Conrad, J.L., Head, J.J., Varricchio, D.J. & Wilson, G.P. (2017) A new Late Cretaceous iguanomorph from North America and the origin of New World Pleurodonta (Squamata, Iguania). *Proceedings of the Royal Society B: Biological Sciences*, **284**, 20161902.

Díez Díaz, V., Garcia, G., Pereda-Suberbiola, X., Jentgen-Ceschino, B., Stein, K., Godefroit, P., *et al.* (2018) The titanosaurian dinosaur Atsinganosaurus velauciensis (Sauropoda) from the Upper Cretaceous of southern France: New material, phylogenetic affinities, and palaeobiogeographical implications. *Cretaceous Research*, **91**, 429–456.

Dong, L., Matsumoto, R., Kusuhashi, N., Wang, Y., Wang, Y. & Evans, S.E. (2020) A new choristodere (Reptilia: Choristodera) from an Aptian–Albian coal deposit in China. *Journal of Systematic Palaeontology*, **18**, 1223–1242.

Dong, L., Roček, Z., Wang, Y. & Jones, M.E.H. (2013) Anurans from the Lower Cretaceous Jehol Group of Western Liaoning, China. *PLOS ONE*, **8**, e69723.

Dong, L., Wang, Y. & Evans, S.E. (2023) A new fossil lizard (Reptilia: Squamata) from the Lower Cretaceous of eastern Inner Mongolia, China. *Cretaceous Research*, **141**, 105363.

Druckenmiller, P.S., Kelley, N.P., Metz, E.T. & Baichtal, J. (2020) An articulated Late Triassic (Norian) thalattosauroid from Alaska and ecomorphology and extinction of Thalattosauria. *Scientific Reports*, **10**, 1746.

Drumheller, S.K. & Wilberg, E.W. (2020) A synthetic approach for assessing the interplay of form and function in the crocodyliform snout. *Zoological Journal of the Linnean Society*, **188**, 507–521.

Drymala, S.M. & Zanno, L.E. (2016) Osteology of *Carnufex carolinensis* (Archosauria: Psuedosuchia) from the Pekin Formation of North Carolina and Its Implications for Early Crocodylomorph Evolution. *PLOS ONE*, **11**, e0157528.

Eltink, E., Da-Rosa, Á.A.S. & Dias-da-Silva, S. (2017) A capitosauroid from the Lower Triassic of South America (Sanga do Cabral Supersequence: Paraná Basin), its phylogenetic relationships and biostratigraphic implications. *Historical Biology*, **29**, 863–874.

Eltink, E. & Dias, E.V. (2012) Temnospôndilos do Brasil: uma breve revisão e aspectos paleobiogeográficos. In *Paleontologia de Vertebrados: Relações entre América do Sul e África. Rio de Janeiro* (ed. by Gallo, V., De Figueiredo, F.J. & Salgado de Carvalho, M.S.). Interciência, pp. 69–98.

Eltink, E., Dias, E.V., Dias-da-Silva, S., Schultz, C.L. & Langer, M.C. (2016) The cranial morphology of the temnospondyl *Australerpeton cosgriffi* (Tetrapoda: Stereospondyli) from the Middle-Late Permian of Paraná Basin and the phylogenetic relationships of Rhinesuchidae. *Zoological Journal of the Linnean Society*, **176**, 835–860.

Eltink, E., Schoch, R.R. & Langer, M.C. (2019) Interrelationships, palaeobiogeography and early evolution of Stereospondylomorpha (Tetrapoda: Temnospondyli). *Journal of Iberian Geology*, **45**, 251–267.

Evans, S. & Matsumoto, R. (2015) An assemblage of lizards from the Early Cretaceous of Japan. *Palaeontologia Electronica*.

Evans, S.E. & Sigogneau-Russell, D. (2001) A stem-group caecilian (Lissamphibia: Gymnophiona) from the Lower Cretaceous of North Africa. *Palaeontology*, **44**, 259–273.

Evers, S.W. & Benson, R.B.J. (2019) A new phylogenetic hypothesis of turtles with implications for the timing and number of evolutionary transitions to marine lifestyles in the group. *Palaeontology*, **62**, 93–134.

Ezcurra, M.D. (2010) A new early dinosaur (Saurischia: Sauropodomorpha) from the Late Triassic of Argentina: a reassessment of dinosaur origin and phylogeny. *Journal of Systematic Palaeontology*, **8**, 371–425.

Ezcurra, M.D. (2016) The phylogenetic relationships of basal archosauromorphs, with an emphasis on the systematics of proterosuchian archosauriforms. *PeerJ*, **4**, e1778.

Ezcurra, M.D., Fiorelli, L.E., Trotteyn, M.J., Martinelli, A.G. & Desojo, J.B. (2020a) The rhynchosaur record, including a new stenaulorhynchine taxon, from the Chañares Formation (upper Ladinian–?lowermost Carnian levels) of La Rioja Province, north-western Argentina. *Journal of Systematic Palaeontology*, **18**, 1907–1938.

Ezcurra, M.D., Nesbitt, S.J., Bronzati, M., Dalla Vecchia, F.M., Agnolin, F.L., Benson, R.B.J., *et al.* (2020b) Enigmatic dinosaur precursors bridge the gap to the origin of Pterosauria. *Nature*, **588**, 445–449.

Ezcurra, M.D., Scheyer, T.M. & Butler, R.J. (2014) The Origin and Early Evolution of Sauria: Reassessing the Permian Saurian Fossil Record and the Timing of the Crocodile-Lizard Divergence. *PLOS ONE*, **9**, e89165.

Fernández, M.S., Campos, L., Maxwell, E.E. & Garrido, A.C. (2021) *Catutosaurus gaspariniae* , gen. et sp. nov. (Ichthyosauria, Thunnosauria) of the Upper Jurassic of Patagonia and the evolution of the ophthalmosaurids. *Journal of Vertebrate Paleontology*, **41**, e1922427.

Fernández-Coll, M., Arbez, T., Bernardini, F. & Fortuny, J. (2019) Cranial anatomy of the Early Triassic trematosaurine *Angusaurus* (Temnospondyli: Stereospondyli): 3D endocranial insights and phylogenetic implications. *Journal of Iberian Geology*, **45**, 269–286.

Field, D.J., Benito, J., Chen, A., Jagt, J.W.M. & Ksepka, D.T. (2020) Late Cretaceous neornithine from Europe illuminates the origins of crown birds. *Nature*, **579**, 397–401.

Flora, H.M. (2019) *A Genus-level Phylogenetic Analysis of Antilocapridae and Implications for the Evolution of Headgear Morphology and Paleoecology* (M.S.).

Foffa, D., Butler, R.J., Nesbitt, S.J., Walsh, S., Barrett, P.M., Brusatte, S.L., *et al.* (2020) Revision of *Erpetosuchus* (Archosauria: Pseudosuchia) and new erpetosuchid material from the Late Triassic 'Elgin Reptile' fauna based on µCT scanning techniques. *Earth and Environmental Science Transactions of the Royal Society of Edinburgh*, **111**, 209–233.

Forasiepi, A.M., Judith Babot, M. & Zimicz, N. (2015) *Australohyaena antiqua* (Mammalia, Metatheria, Sparassodonta), a large predator from the Late Oligocene of Patagonia. *Journal of Systematic Palaeontology*, **13**, 503–525.

Ford, D.P. & Benson, R.B.J. (2020) The phylogeny of early amniotes and the affinities of Parareptilia and Varanopidae. *Nature Ecology & Evolution*, **4**, 57–65.

Foth, C. & Joyce, W.G. (2016) Slow and steady: the evolution of cranial disparity in fossil and recent turtles. *Proceedings of the Royal Society B: Biological Sciences*, **283**, 20161881.

Fraser-King, S.W., Benoit, J., Day, M.O. & Rubidge, B.S. (2019) Cranial morphology and phylogenetic relationship of the enigmatic dinocephalian *Styracocephalus platyrhynchus* from the Karoo Supergroup, South Africa. *Palaeontologia Africana*, **54**, 14–29.

Funston, G.F., Chinzorig, T., Tsogtbaatar, K., Kobayashi, Y., Sullivan, C. & Currie, P.J. (2020) A new two-fingered dinosaur sheds light on the radiation of Oviraptorosauria. *Royal Society Open Science*, **7**, 201184.

Funston, G.F. & Currie, P.J. (2016) A new caenagnathid (Dinosauria: Oviraptorosauria) from the Horseshoe Canyon Formation of Alberta, Canada, and a reevaluation of the relationships of Caenagnathidae. *Journal of Vertebrate Paleontology*, **36**, e1160910.

Gaetán, C.M., Buono, M.R. & Gaetano, L.C. (2018) *Prosqualodon australis* (Cetacea: Odontoceti) from the Early Miocene of Patagonia, Argentina: Redescription and Phylogenetic Analysis. *Ameghiniana*, **56**, 1–27.

Gallina, P.A. & Apesteguía, S. (2011) Cranial Anatomy and Phylogenetic Position of the Titanosaurian Sauropod Bonitasaura salgadoi. *Acta Palaeontologica Polonica*, **56**, 45–60.

Gao, K.-Q. & Chen, J. (2017) A New Crown-Group Frog (Amphibia: Anura) from the Early Cretaceous of Northeastern Inner Mongolia, China. *American Museum Novitates*, **2017**, 1–39.

Gao, K.-Q. & Shubin, N.H. (2001) Late Jurassic salamanders from northern China. *Nature*, **410**, 574–577.

Gao, K.-Q. & Shubin, N.H. (2012) Late Jurassic salamandroid from western Liaoning, China. *Proceedings of the National Academy of Sciences*, **109**, 5767–5772.

Gardner, J.D. (2003) Revision of *Habrosaurus* Gilmore (Caudata; Sirenidae) and relationships among sirenid salamanders. *Palaeontology*, **46**, 1089–1122.

Gee, B.M. (2020) Size matters: the effects of ontogenetic disparity on the phylogeny of Trematopidae (Amphibia: Temnospondyli). *Zoological Journal of the Linnean Society*, **190**, 79–113.

Gee, B.M. (2021) Returning to the roots: resolution, reproducibility, and robusticity in the phylogenetic inference of Dissorophidae (Amphibia: Temnospondyli). *PeerJ*, **9**, e12423.

Gee, B.M., Parker, W.G. & Marsh, A.D. (2020) Redescription of *Anaschisma* (Temnospondyli: Metoposauridae) from the Late Triassic of Wyoming and the phylogeny of the Metoposauridae. *Journal of Systematic Palaeontology*, **18**, 233–258.

Geisler, J.H. (2001) New Morphological Evidence for the Phylogeny of Artiodactyla, Cetacea, and Mesonychidae. *American Museum Novitates*, **2001**, 1–53.

Gentry, A.D. (2016) New material of the Late Cretaceous marine turtle Ctenochelys acris Zangerl, 1953 and a phylogenetic reassessment of the 'toxochelyid'-grade taxa. *Journal of Systematic Palaeontology*, **15**, 675–696.

Gentry, A.D., Kiernan, C.R. & Parham, J.F. (2023) A large non-marine turtle from the Upper Cretaceous of Alabama and a review of North American "Macrobaenids." *The Anatomical Record*, **306**, 1411–1430.

Germain, D. (2010) The Moroccan diplocaulid: the last lepospondyl, the single one on Gondwana. *Historical Biology*, **22**, 4–39.

Geroto, C.F.C. & Bertini, R.J. (2019) New material of *Pepesuchus* (Crocodyliformes; Mesoeucrocodylia) from the Bauru Group: implications about its phylogeny and the age of the Adamantina Formation. *Zoological Journal of the Linnean Society*, **185**, 312–334.

Gess, R. & Ahlberg, P.E. (2018) A tetrapod fauna from within the Devonian Antarctic Circle. *Science*, **360**, 1120–1124.

Gheerbrant, E., Khaldoune, F., Schmitt, A. & Tabuce, R. (2021) Earliest Embrithopod Mammals (Afrotheria, Tethytheria) from the Early Eocene of Morocco: Anatomy, Systematics and Phylogenetic Significance. *Journal of Mammalian Evolution*, **28**, 245–283.

Gianechini, F.A., Méndez, A.H., Filippi, L.S., Paulina-Carabajal, A., Juárez-Valieri, R.D. & Garrido, A.C. (2020) A new furileusaurian abelisaurid from La Invernada (Upper Cretaceous, Santonian, Bajo de la Carpa Formation), northern Patagonia, Argentina. *Journal of Vertebrate Paleontology*, **40**, e1877151.

Gorscak, E., O'Connor, P.M., Roberts, E.M. & Stevens, N.J. (2017) The second titanosaurian (Dinosauria: Sauropoda) from the middle Cretaceous Galula Formation, southwestern Tanzania, with remarks on African titanosaurian diversity. *Journal of Vertebrate Paleontology*, **37**, e1343250.

Grealy, A., Phillips, M., Miller, G., Gilbert, M.T.P., Rouillard, J.-M., Lambert, D., *et al.* (2017) Eggshell palaeogenomics: Palaeognath evolutionary history revealed through ancient nuclear and mitochondrial DNA from Madagascan elephant bird (Aepyornis sp.) eggshell. *Molecular Phylogenetics and Evolution*, **109**, 151–163.

Griffiths, E.F., Ford, D.P., Benson, R.B.J. & Evans, S.E. (2021) New information on the Jurassic lepidosauromorph Marmoretta oxoniensis. *Papers in Palaeontology*, **7**, 2255–2278.

Groh, S.S., Upchurch, P., Barrett, P.M. & Day, J.J. (2020) The phylogenetic relationships of neosuchian crocodiles and their implications for the convergent evolution of the longirostrine condition. *Zoological Journal of the Linnean Society*, **188**, 473–506.

Hamley, T., Cisneros, J.C. & Damiani, R. (2021) A procolophonid reptile from the Lower Triassic of Australia. *Zoological Journal of the Linnean Society*, **192**, 554–609.

Hartman, S., Mortimer, M., Wahl, W.R., Lomax, D.R., Lippincott, J. & Lovelace, D.M. (2019) A new paravian dinosaur from the Late Jurassic of North America supports a late acquisition of avian flight. *PeerJ*, **7**, e7247.

Heckert, A.B., Nesbitt, S.J., Stocker, M.R., Schneider, V.P., Hoffman, D.K. & Zimmer, B.W. (2021) A new short-faced archosauriform from the Upper Triassic Placerias/Downs' quarry complex, Arizona, USA, expands the morphological diversity of the Triassic archosauriform radiation. *The Science of Nature*, **108**, 32.

Heintzman, P.D., Zazula, G.D., Cahill, J.A., Reyes, A.V., MacPhee, R.D.E. & Shapiro, B. (2015) Genomic Data from Extinct North American Camelops Revise Camel Evolutionary History. *Molecular Biology and Evolution*, **32**, 2433–2440.

Hendrickx, C., Hartman, S. & Mateus, O. (2015) An overview of non-avian theropod discoveries and classification. *PalArch's Journal of Vertebrate Palaeontology*, **12**, 1–73.

Henrici, A.C., Báez, A.M. & Grande, L. (2013) *Aerugoamnis paulus*, New Genus and New Speices (Anura: Anomocoela): First Reported Anuran from the Early Eocene (Wasatchian) Fossil Butte Member of the Green River Formation, Wyoming. *Annals of Carnegie Museum*, **81**, 295–309.

Herrera, J.P. & Dávalos, L.M. (2016) Phylogeny and Divergence Times of Lemurs Inferred with Recent and Ancient Fossils in the Tree. *Systematic Biology*, **65**, 772–791.

Herrera-Flores, J.A., Stubbs, T.L. & Benton, M.J. (2017) Macroevolutionary patterns in Rhynchocephalia: is the tuatara (Sphenodon punctatus) a living fossil? *Palaeontology*, **60**, 319–328.

Hirayama, R. (1997) Chapter 8 - Distribution and Diversity of Cretaceous Chelonioids. In *Ancient Marine Reptiles* (ed. by Callaway, J.M. & Nicholls, E.L.). Academic Press, San Diego, pp. 225–241.

Holley, J.A., Sterli, J. & Basso, N.G. (2020) Dating the origin and diversification of Pan-Chelidae (Testudines, Pleurodira) under multiple molecular clock approaches. *Contributions to Zoology*, **89**, 146–174.

Hsiou, A.S., Nydam, R.L., Simões, T.R., Pretto, F.A., Onary, S., Martinelli, A.G., *et al.* (2019) A New Clevosaurid from the Triassic (Carnian) of Brazil and the Rise of Sphenodontians in Gondwana. *Scientific Reports*, **9**, 11821.

Huang, Y.N., Li, J., Jiang, Q.Y., Shen, X.S., Yan, X.Y., Tang, Y.B., *et al.* (2015) Complete mitochondrial genome of the *Cyclemys dentata* and phylogenetic analysis of the major family Geoemydidae. *Genetics and molecular research: GMR*, **14**, 3234–3243.

Huebinger, R., Bickham, J., Rhodin, A. & Mittermeier, R. (2013) Mitochondrial DNA Corroborates Taxonomy of the South American Chelid Turtles of the Genera Platemys and Acanthochelys. *Chelonian Conservation and Biology*, **12**, 168–171.

Hungerbühler, A. (2001) The status and phylogenetic relationships of *"Zanclodon"arenaceus*: the earliest known phytosaur? *Paläontologische Zeitschrift*, **75**, 97–112.

Huttenlocker, A.K., Pardo, J.D., Small, B.J. & Anderson, J.S. (2013) Cranial morphology of recumbirostrans (Lepospondyli) from the Permian of Kansas and Nebraska, and early morphological evolution inferred by micro-computed tomography. *Journal of Vertebrate Paleontology*, **33**, 540–552.

Huttenlocker, A.K. & Sidor, C.A. (2020) A Basal Nonmammaliaform Cynodont from the Permian of Zambia and the Origins of Mammalian Endocranial and Postcranial Anatomy. *Journal of Vertebrate Paleontology*, **40**, e1827413.

Huttenlocker, A.K., Singh, S.A., Henrici, A.C. & Sumida, S.S. (2021) A Carboniferous synapsid with caniniform teeth and a reappraisal of mandibular size-shape heterodonty in the origin of mammals. *Royal Society Open Science*, **8**, 211237.

Ibiricu, L.M., Baiano, M.A., Martínez, R.D., Alvarez, B.N., Lamanna, M.C. & Casal, G.A. (2021) A detailed osteological description of Xenotarsosaurus bonapartei (Theropoda: Abelisauridae): implications for abelisauroid phylogeny. *Cretaceous Research*, **124**, 104829.

Ikeda, T., Ota, H. & Matsui, M. (2016) New fossil anurans from the Lower Cretaceous Sasayama Group of Hyogo Prefecture, Western Honshu, Japan. *Cretaceous Research*, **61**, 108–123.

Ikeda, T., Ota, H. & Saegusa, H. (2015) A new fossil lizard from the Lower Cretaceous Sasayama Group of Hyogo Prefecture, western Honshu, Japan. *Journal of Vertebrate Paleontology*, **35**, e885032.

Imai, T., Azuma, Y., Kawabe, S., Shibata, M., Miyata, K., Wang, M., *et al.* (2019) An unusual bird (Theropoda, Avialae) from the Early Cretaceous of Japan suggests complex evolutionary history of basal birds. *Communications Biology*, **2**, 1–11.

Ivakhnenko, M.F. (2012) Permian Cynodontia (Theromorpha) of Eastern Europe. *Paleontological Journal*, **46**, 199–207.

Jackson, D.R. (1988) A Re-Examination of Fossil Turtles of the Genus Trachemys (Testudines: Emydidae). *Herpetologica*, **44**, 317–325.

Jetz, W., Thomas, G.H., Joy, J.B., Hartmann, K. & Mooers, A.O. (2012) The global diversity of birds in space and time. *Nature*, **491**, 444–448.

Jia, J. & Gao, K.-Q. (2016) A New Basal Salamandroid (Amphibia, Urodela) from the Late Jurassic of Qinglong, Hebei Province, China. *PLOS ONE*, **11**, e0153834.

Jiang, D.-Y., Hao, W.-C., Maisch, M.W., Matzke, A.T. & Sun, Y.-L. (2005) A Basal Mixosaurid Ichthyosaur from the Middle Triassic of China. *Palaeontology*, **48**, 869–882.

Jiangzuo, Q., Werdelin, L. & Sun, Y. (2022) A dwarf sabertooth cat (Felidae: Machairodontinae) from Shanxi, China, and the phylogeny of the sabertooth tribe Machairodontini. *Quaternary Science Reviews*, **284**, 107517.

Johnson, M.M., Young, M.T. & Brusatte, S.L. (2020) The phylogenetics of Teleosauroidea (Crocodylomorpha, Thalattosuchia) and implications for their ecology and evolution. *PeerJ*, **8**, e9808.

Jones, A.S. & Butler, R.J. (2018) A new phylogenetic analysis of Phytosauria (Archosauria: Pseudosuchia) with the application of continuous and geometric morphometric character coding. *PeerJ*, **6**, e5901.

Jones, M.E., Anderson, C.L., Hipsley, C.A., Müller, J., Evans, S.E. & Schoch, R.R. (2013) Integration of molecules and new fossils supports a Triassic origin for Lepidosauria (lizards, snakes, and tuatara). *BMC Evolutionary Biology*, **13**, 208.

Joyce, W.G., Lyson, T.R. & Kirkland, J.I. (2016) An early bothremydid (Testudines, Pleurodira) from the Late Cretaceous (Cenomanian) of Utah, North America. *PeerJ*, **4**, e2502.

Kear, B.P., Lee, M.S.Y., Gerdtz, W.R. & Flannery, T.F. (2008) Evolution of Hind Limb Proportions in Kangaroos (Marsupialia: Macropodoidea). In *Mammalian Evolutionary*

*Morphology: A Tribute to Frederick S. Szalay*, Vertebrate Paleobiology and Paleoanthropology Series (ed. by Sargis, E.J. & Dagosto, M.). Springer Netherlands, Dordrecht, pp. 25–35.

Kelly, T.S. & Martin, R.A. (2022) Phylogenetic positions of *Paronychomys* Jacobs and *Basirepomys* Korth and De Blieux relative to the tribe Neotomini (Rodentia, Cricetidae). *Journal of Paleontology*, **96**, 692–705.

Kerber, L., Mayer, E.L., Gomes, A.C. & Nasif, N. (2020) On the morphological, taxonomic, and phylogenetic status of South American Quaternary dinomyid rodents (Rodentia: Dinomyidae). *PalZ*, **94**, 167–178.

Klein, N., Furrer, H., Ehrbar, I., Torres Ladeira, M., Richter, H. & Scheyer, T.M. (2022) A new pachypleurosaur from the Early Ladinian Prosanto Formation in the Eastern Alps of Switzerland. *Swiss Journal of Palaeontology*, **141**, 12.

Kligman, B.T., Gee, B.M., Marsh, A.D., Nesbitt, S.J., Smith, M.E., Parker, W.G., *et al.* (2023) Triassic stem caecilian supports dissorophoid origin of living amphibians. *Nature*, **614**, 102–107.

Kurkin, A.A. (2017) A new Galeopid (Anomodontia, Galeopidae) from the Permian of Eastern Europe. *Paleontological Journal*, **51**, 308–312.

Kutty, T.S., Chatterjee, S., Galton, P.M. & Upchurch, P. (2007) Basal Sauropodomorphs (Dinosauria: Saurischia) From The Lower Jurassic Of India: Their Anatomy And Relationships. *Journal of Paleontology*, **81**, 1218–1240.

Laboury, A., Bennion, R.F., Thuy, B., Weis, R. & Fischer, V. (2022) Anatomy and phylogenetic relationships of *Temnodontosaurus zetlandicus* (Reptilia: Ichthyosauria). *Zoological Journal of the Linnean Society*, **195**, 172–194.

Lacerda, M.B., França, M.A.G. de & Schultz, C.L. (2018) A new erpetosuchid (Pseudosuchia, Archosauria) from the Middle–Late Triassic of Southern Brazil. *Zoological Journal of the Linnean Society*, **184**, 804–824.

Laloy, F., Rage, J.-C., Evans, S.E., Boistel, R., Lenoir, N. & Laurin, M. (2013) A Re-Interpretation of the Eocene Anuran *Thaumastosaurus* Based on MicroCT Examination of a 'Mummified' Specimen. *PLOS ONE*, **8**, e74874.

Langer, M.C., Ezcurra, M.D., Rauhut, O.W.M., Benton, M.J., Knoll, F., McPhee, B.W., *et al.* (2017) Untangling the dinosaur family tree. *Nature*, **551**, E1–E3.

Laurin, M. & Soler-Gijón, R. (2001) The oldest stegocephalian from the Iberian Peninsula: evidence that temnospondyls were euryhaline. *Comptes Rendus de l'Académie des Sciences - Series III - Sciences de la Vie*, **324**, 495–501.

Le, M., Raxworthy, C.J., McCord, W.P. & Mertz, L. (2006) A molecular phylogeny of tortoises (Testudines: Testudinidae) based on mitochondrial and nuclear genes. *Molecular Phylogenetics and Evolution*, **40**, 517–531.

Lecuona, A., Ezcurra, M.D. & Irmis, R.B. (2016) Revision of the early crocodylomorph *Trialestes romeri* (Archosauria, Suchia) from the lower Upper Triassic Ischigualasto Formation of Argentina: one of the oldest-known crocodylomorphs. *Papers in Palaeontology*, **2**, 585–622.

Lee, M.S.Y., Caldwell, M.W. & Scanlon, J.D. (1999) A second primitive marine snake: *Pachyophis woodwardi* from the Cretaceous of Bosnia-Herzegovina. *Journal of Zoology*, **248**, 509–520.

Lee, S., Lee, Y.-N., Chinsamy, A., Lü, J., Barsbold, R. & Tsogtbaatar, K. (2019) A new baby oviraptorid dinosaur (Dinosauria: Theropoda) from the Upper Cretaceous Nemegt Formation of Mongolia. *PLOS ONE*, **14**, e0210867.

Li, C., Fraser, N.C., Rieppel, O. & Wu, X.-C. (2018) A Triassic stem turtle with an edentulous beak. *Nature*, **560**, 476–479.

Li, Q. & Liu, J. (2020) An Early Triassic sauropterygian and associated fauna from South China provide insights into Triassic ecosystem health. *Communications Biology*, **3**, 1–11.

Liu, J. & Abdala, N.F. (2023) Late Permian terrestrial faunal connections invigorated: the first whaitsioid therocephalian from China. *Palaeontologia Africana*, **56**, 111–117.

Liu, J. & Chen, J. (2020) The tetrapod fauna of the upper Permian Naobaogou Formation of China: 7. *Laosuchus hun* sp. nov. (Chroniosuchia) and interrelationships of chroniosuchians. *Journal of Systematic Palaeontology*, **18**, 2043–2058.

Longrich, N.R., Bardet, N., Khaldoune, F., Yazami, O.K. & Jalil, N.-E. (2021) *Pluridens serpentis*, a new mosasaurid (Mosasauridae: Halisaurinae) from the Maastrichtian of Morocco and implications for mosasaur diversity. *Cretaceous Research*, **126**, 104882.

Longrich, N.R., Pereda-Suberbiola, X., Jalil, N.-E., Khaldoune, F. & Jourani, E. (2017) An abelisaurid from the latest Cretaceous (late Maastrichtian) of Morocco, North Africa. *Cretaceous Research*, **76**, 40–52.

Longrich, N.R., Vinther, J., Pyron, R.A., Pisani, D. & Gauthier, J.A. (2015) Biogeography of worm lizards (Amphisbaenia) driven by end-Cretaceous mass extinction. *Proceedings of the Royal Society B: Biological Sciences*, **282**, 20143034.

Louchart, A. & Viriot, L. (2011) From snout to beak: the loss of teeth in birds. *Trends in Ecology & Evolution*, **26**, 663–673.

Lucas, S.G. & Oakes, W. (1988) A Late Triassic Cynodont from the American South-West. *Palaeontology*, **31**, 445–449.

Luo, Z.-X., Yuan, C.-X., Meng, Q.-J. & Ji, Q. (2011) A Jurassic eutherian mammal and divergence of marsupials and placentals. *Nature*, **476**, 442–445.

Lyson, T.R., Joyce, W.G., Knauss, G.E. & Pearson, D.A. (2011) *Boremys* (Testudines, Baenidae) from the latest Cretaceous and early Paleocene of North Dakota: an 11-millionyear range extension and an additional K/T survivor. *Journal of Vertebrate Paleontology*, **31**, 729–737.

Macdonald, I. & Currie, P.J. (2019) Description of a partial *Dromiceiomimus* (Dinosauria: Theropoda) skeleton with comments on the validity of the genus. *Canadian Journal of Earth Sciences*, **56**, 129–157.

Maddin, H.C., Jenkins Jr., F.A. & Anderson, J.S. (2012) The Braincase of *Eocaecilia micropodia* (Lissamphibia, Gymnophiona) and the Origin of Caecilians. *PLOS ONE*, **7**, e50743.

Maisch, M.W. (2017) Re-assessment of *Silphoictidoides ruhuhuensis* von Huene, 1950 (Therapsida, Therocephalia) from the Late Permian of Tanzania: one of the most basal baurioids known. *Palaeodiversity*, **10**, 25–39.

Mallon, J.C. & Brinkman, D.B. (2018) *Basilemys morrinensis* , a new species of nanhsiungchelyid turtle from the Horseshoe Canyon Formation (Upper Cretaceous) of Alberta, Canada. *Journal of Vertebrate Paleontology*, **38**, e1431922.

Mann, A., Calthorpe, A.S. & Maddin, H.C. (2021) *Joermungandr bolti* , an exceptionally preserved 'microsaur' from the Mazon Creek Lagerstätte reveals patterns of integumentary evolution in Recumbirostra. *Royal Society Open Science*, **8**, 210319.

Mann, A. & Maddin, H.C. (2019) *Diabloroter bolti*, a short-bodied recumbirostran 'microsaur' from the Francis Creek Shale, Mazon Creek, Illinois. *Zoological Journal of the Linnean Society*, **187**, 494–505.

Mann, A., Pardo, J.D. & Maddin, H.C. (2019) *Infernovenator steenae*, a new serpentine recumbirostran from the 'Mazon Creek' *Lagerstätte* further clarifies lysorophian origins. *Zoological Journal of the Linnean Society*, **187**, 506–517.

Mannion, P.D., Upchurch, P., Schwarz, D. & Wings, O. (2019) Taxonomic affinities of the putative titanosaurs from the Late Jurassic Tendaguru Formation of Tanzania: phylogenetic and biogeographic implications for eusauropod dinosaur evolution. *Zoological Journal of the Linnean Society*, **185**, 784–909.

Marjanović, D. & Laurin, M. (2014) An updated paleontological timetree of lissamphibians, with comments on the anatomy of Jurassic crown-group salamanders (Urodela). *Historical Biology*, **26**, 535–550.

Marsicano, C., Angielczyk, K.D., Cisneros, J.C., Richter, M., Kammerer, C.F., Fröbisch, J., *et al.* (2021) Brazilian Permian Dvinosaurs (Amphibia, Temnospondyli): Revised Description and Phylogeny. *Journal of Vertebrate Paleontology*, e1893181.

Marsicano, C.A., Latimer, E., Rubidge, B. & Smith, R.M.H. (2017) The Rhinesuchidae and early history of the Stereospondyli (Amphibia: Temnospondyli) at the end of the Palaeozoic. *Zoological Journal of the Linnean Society*, **181**, 357–384.

Martínez, R.D.F., Lamanna, M.C., Novas, F.E., Ridgely, R.C., Casal, G.A., Martínez, J.E., *et al.* (2016) A Basal Lithostrotian Titanosaur (Dinosauria: Sauropoda) with a Complete Skull: Implications for the Evolution and Paleobiology of Titanosauria. *PLOS ONE*, **11**, e0151661.

Marx, M.P., Mateus, O., Polcyn, M.J., Schulp, A.S., Gonçalves, A.O. & Jacobs, L.L. (2021) The cranial anatomy and relationships of *Cardiocorax mukulu* (Plesiosauria: Elasmosauridae) from Bentiaba, Angola. *PLOS ONE*, **16**, e0255773.

Marzola, M., Mateus, O., Shubin, N.H. & Clemmensen, L.B. (2017) *Cyclotosaurus naraserluki,* sp. nov., a new Late Triassic cyclotosaurid (Amphibia, Temnospondyli) from the Fleming Fjord Formation of the Jameson Land Basin (East Greenland). *Journal of Vertebrate Paleontology*, **37**, e1303501.

Matamales-Andreu, R., Roig-Munar, F.X., Oms, O., Galobart, À. & Fortuny, J. (2021) A captorhinid-dominated assemblage from the palaeoequatorial Permian of Menorca (Balearic Islands, western Mediterranean). *Earth and Environmental Science Transactions of The Royal Society of Edinburgh*, **112**, 125–145.

Mateus, O. & Estraviz-López, D. (2022) A new theropod dinosaur from the early cretaceous (Barremian) of Cabo Espichel, Portugal: Implications for spinosaurid evolution. *PLOS ONE*, **17**, e0262614.

Matsumoto, R., Dong, L., Wang, Y. & Evans, S.E. (2019) The first record of a nearly complete choristodere (Reptilia: Diapsida) from the Upper Jurassic of Hebei Province, People's Republic of China. *Journal of Systematic Palaeontology*, **17**, 1031–1048.

Matsumoto, R. & Evans, S.E. (2018) The first record of albanerpetontid amphibians (Amphibia: Albanerpetontidae) from East Asia. *PLOS ONE*, **13**, e0189767.

Mattingly, S.G., Beard, K.C., Coster, P.M.C., Salem, M.J., Chaimanee, Y. & Jaeger, J.-J. (2021) A new parapithecine (Primates: Anthropoidea) from the early Oligocene of Libya supports parallel evolution of large body size among parapithecids. *Journal of Human Evolution*, **153**, 102957.

Maxwell, E.E., Cortés, D., Patarroyo, P. & Ruge, M.L.P. (2019) A new specimen of Platypterygius sachicarum (Reptilia, Ichthyosauria) from the Early Cretaceous of Colombia and its phylogenetic implications. *Journal of Vertebrate Paleontology*, **39**, e1577875.

Mayr, G. (2020) A remarkably complete skeleton from the London Clay provides insights into the morphology and diversity of early Eocene zygodactyl near-passerine birds. *Journal of Systematic Palaeontology*, **18**, 1891–1906.

McFeeters, B., Ryan, M.J., Schröder-Adams, C. & Cullen, T.M. (2016) A new ornithomimid theropod from the Dinosaur Park Formation of Alberta, Canada. *Journal of Vertebrate Paleontology*, **36**, e1221415.

Menegazzo, M.C., Bertini, R.J. & Manzini, F.F. (2015) A new turtle from the Upper Cretaceous Bauru Group of Brazil, updated phylogeny and implications for age of the Santo Anastácio Formation. *Journal of South American Earth Sciences*, **58**, 18–32.

Meng, Y., Da-Qing, L.I., Ksepka, D.T. & Hong-Yu, Y.I. (2021) A juvenile skull of the longirostrine choristodere (Diapsida: Choristodera), *Mengshanosaurus minimus* gen. et sp. nov., with comments on neochoristodere ontogeny. *Vertebrata Palasiatica*, **59**, 213.

Miguel Chaves, C. de, Ortega, F. & Pérez-García, A. (2018) New highly pachyostotic nothosauroid interpreted as a filter-feeding Triassic marine reptile. *Biology Letters*, **14**, 20180130.

Mihlbachler, M.C. (2008) Species Taxonomy, Phylogeny, and Biogeography of the Brontotheriidae (Mammalia: Perissodactyla). *Bulletin of the American Museum of Natural History*, **2008**, 1–475.

Milner, A. & Schoch, R. (2013) *Trimerorhachis* (Amphibia: Temnospondyli) from the Lower Permian of Texas and New Mexico: Cranial osteology, taxonomy and biostratigraphy. *Neues Jahrbuch für Geologie und Paläontologie - Abhandlungen*, **270**, 91–128.

Milner, A.C., Milner, A.R. & Walsh, S.A. (2009) A new specimen of *Baphetes* from Nýřany, Czech Republic and the intrinsic relationships of the Baphetidae. *Acta Zoologica*, **90**, 318–334.

Modesto, S.P. & Damiani, R. (2007) The procolophonoid reptile *Sauropareion anoplus* from the lowermost Triassic of South Africa. *Journal of Vertebrate Paleontology*, **27**, 337–349.

Modesto, S.P., Damiani, R.J., Neveling, J. & Yates, A.M. (2003) A new Triassic owenettid parareptile and the Mother of Mass Extinctions. *Journal of Vertebrate Paleontology*, **23**, 715–719.

Modesto, S.P., Richards, C.D., Ide, O. & Sidor, C.A. (2018) The vertebrate fauna of the Upper Permian of Niger—X. The mandible of the captorhinid reptile *Moradisaurus grandis*. *Journal of Vertebrate Paleontology*, **38**, e1531877.

Moon, B.C. (2019) A new phylogeny of ichthyosaurs (Reptilia: Diapsida). *Journal of Systematic Palaeontology*, **17**, 129–155.

Moore, A.J., Barrett, P.M., Upchurch, P., Liao, C.-C., Ye, Y., Hao, B., *et al.* (2023) Re-assessment of the Late Jurassic eusauropod Mamenchisaurus sinocanadorum Russell and Zheng, 1993, and the evolution of exceptionally long necks in mamenchisaurids. *Journal of Systematic Palaeontology*, **21**, 2171818.

Müller, J., Li, J.-L. & Reisz, R.R. (2008) A new bolosaurid parareptile, *Belebey chengi* sp. nov., from the Middle Permian of China and its paleogeographic significance. *Naturwissenschaften*, **95**, 1169–1174.

Müller, J. & Reisz, R.R. (2006) The Phylogeny of Early Eureptiles: Comparing Parsimony and Bayesian Approaches in the Investigation of a Basal Fossil Clade. *Systematic Biology*, **55**, 503–511.

Müller, R., Langer, M. & Dias-da-Silva, S. (2018) Ingroup relationships of Lagerpetidae (Avemetatarsalia: Dinosauromorpha): A further phylogenetic investigation on the understanding of dinosaur relatives. *Zootaxa*, **4392**, 149.

Muzzopappa, P., Martinelli, A.G., Garderes, J.P. & Rougier, G.W. (2021) Exceptional avian pellet from the Paleocene of Patagonia and description of its content: a new species of calyptocephalellid (Neobatrachia) anuran. *Papers in Palaeontology*, **7**, 1133–1146.

Napoli, J.G., Ruebenstahl, A.A., Bhullar, B.-A.S., Turner, A.H. & Norell, M.A. (2021) A New Dromaeosaurid (Dinosauria: Coelurosauria) from Khulsan, Central Mongolia. *American Museum Novitates*, **2021**, 1–47.

Nesbitt, S.J. (2011) The Early Evolution of Archosaurs: Relationships and the Origin of Major Clades. *Bulletin of the American Museum of Natural History*, **2011**, 1–292.

Nesbitt, S.J., Butler, R.J., Ezcurra, M.D., Barrett, P.M., Stocker, M.R., Angielczyk, K.D., *et al.* (2017) The earliest bird-line archosaurs and the assembly of the dinosaur body plan. *Nature*, **544**, 484–487.

Nesbitt, S.J., Denton, R.K., Loewen, M.A., Brusatte, S.L., Smith, N.D., Turner, A.H., *et al.* (2019) A mid-Cretaceous tyrannosauroid and the origin of North American end-Cretaceous dinosaur assemblages. *Nature Ecology & Evolution*, **3**, 892–899.

Nesbitt, S.J., Stocker, M.R., Ezcurra, M.D., Fraser, N.C., Heckert, A.B., Parker, W.G., *et al.* (2022) Widespread azendohsaurids (Archosauromorpha, Allokotosauria) from the Late Triassic of western USA and India. *Papers in Palaeontology*, **8**, e1413.

Norman, D.B., FLS, Baron, M.G., Garcia, M.S. & Müller, R.T. (2022) Taxonomic, palaeobiological and evolutionary implications of a phylogenetic hypothesis for Ornithischia (Archosauria: Dinosauria). *Zoological Journal of the Linnean Society*, **196**, 1273–1309.

Olive, S., Leroy, Y., Daeschler, E.B., Downs, J.P., Ladevèze, S. & Clément, G. (2020) Tristichopterids (Sarcopterygii, Tetrapodomorpha) from the Upper Devonian tetrapod-bearing locality of Strud (Belgium, upper Famennian), with phylogenetic and paleobiogeographic considerations. *Journal of Vertebrate Paleontology*, **40**, e1768105.

Oriozabala, C., Sterli, J. & Fuente, M.S. de la. (2020) New species of the long-necked chelid Yaminuechelys from the Upper Cretaceous (Campanian–Maastrichtian) of Chubut, Argentina. *Cretaceous Research*, **106**, 104197.

Osmólska, H. (1996) An unusual theropod dinosaur from the Late Cretaceous Nemegt Formation of Mongolia. *Acta Palaeontologica Polonica*, **41**, 1–38.

Otero, A., Krupandan, E., Pol, D., Chinsamy, A. & Choiniere, J. (2015) A new basal sauropodiform from South Africa and the phylogenetic relationships of basal sauropodomorphs. *Zoological Journal of the Linnean Society*, **174**, 589–634.

Otero, A. & Pol, D. (2021) Ontogenetic changes in the postcranial skeleton of *Mussaurus patagonicus* (Dinosauria, Sauropodomorpha) and their impact on the phylogenetic relationships of early sauropodomorphs. *Journal of Systematic Palaeontology*, **19**, 1467–1516.

Pacheco, C., Müller, R.T., Langer, M., Pretto, F.A., Kerber, L. & Silva, S.D. da. (2019) *Gnathovorax cabreirai*: a new early dinosaur and the origin and initial radiation of predatory dinosaurs. *PeerJ*, **7**, e7963.

Pacheco, C.P., Eltink, E., Müller, R.T. & Dias-da-Silva, S. (2017) A new Permian temnospondyl with Russian affinities from South America, the new family Konzhukoviidae, and the phylogenetic status of Archegosauroidea. *Journal of Systematic Palaeontology*, **15**, 241–256.

Panciroli, E., Benson, R.B.J., Fernandez, V., Butler, R.J., Fraser, N.C., Luo, Z.-X., *et al.* (2021) New species of mammaliaform and the cranium of *Borealestes* (Mammaliformes: Docodonta) from the Middle Jurassic of the British Isles. *Zoological Journal of the Linnean Society*, **192**, 1323–1362.

Panciroli, E., Benson, R.B.J. & Walsh, S. (2017) The dentary of *Wareolestes rex* (Megazostrodontidae): a new specimen from Scotland and implications for morganucodontan tooth replacement. *Papers in Palaeontology*, **3**, 373–386.

Páramo-Fonseca, M.E., O'Gorman, J.P., Gasparini, Z., Padilla, S. & Parra-Ruge, M.L. (2019) A new late Aptian elasmosaurid from the Paja Formation, Villa de Leiva, Colombia. *Cretaceous Research*, **99**, 30–40.

Pardo, J.D. & Mann, A. (2018) A basal aïstopod from the earliest Pennsylvanian of Canada, and the antiquity of the first limbless tetrapod lineage. *Royal Society Open Science*, **5**, 181056.

Pardo, J.D., Small, B.J. & Huttenlocker, A.K. (2017a) Stem caecilian from the Triassic of Colorado sheds light on the origins of Lissamphibia. *Proceedings of the National Academy of Sciences*, **114**, E5389–E5395.

Pardo, J.D., Szostakiwskyj, M., Ahlberg, P.E. & Anderson, J.S. (2017b) Hidden morphological diversity among early tetrapods. *Nature*, **546**, 642–645.

Parisi Dutra, R., Casali, D. de M., Missagia, R.V., Gasparini, G.M., Perini, F.A. & Cozzuol, M.A. (2017) Phylogenetic Systematics of Peccaries (Tayassuidae: Artiodactyla) and a Classification of South American Tayassuids. *Journal of Mammalian Evolution*, **24**, 345–358.

Patrick, E.L., Whiteside, D.I. & Benton, M.J. (2019) A new crurotarsan archosaur from the Late Triassic of South Wales. *Journal of Vertebrate Paleontology*, **39**, e1645147.

Pattinson, D.J., Thompson, R.S., Piotrowski, A.K. & Asher, R.J. (2015) Phylogeny, Paleontology, and Primates: Do Incomplete Fossils Bias the Tree of Life? *Systematic Biology*, **64**, 169–186.

Pêgas, R.V., Costa, F.R. & Kellner, A.W.A. (2018) New Information on the osteology and a taxonomic revision of The genus Thalassodromeus (Pterodactyloidea, Tapejaridae, Thalassodrominae). *Journal of Vertebrate Paleontology*, **38**, e1443273.

Pêgas, R.V., Holgado, B., Ortiz David, L.D., Baiano, M.A. & Costa, F.R. (2022) On the pterosaur *Aerotitan sudamericanus* (Neuquén Basin, Upper Cretaceous of Argentina), with comments on azhdarchoid phylogeny and jaw anatomy. *Cretaceous Research*, **129**, 104998.

Pei, R., Pittman, M., Goloboff, P.A., Dececchi, T.A., Habib, M.B., Kaye, T.G., *et al.* (2020) Potential for Powered Flight Neared by Most Close Avialan Relatives, but Few Crossed Its Thresholds. *Current Biology*, **30**, 4033-4046.e8.

Pei, R., Qin, Y., Wen, A., Zhao, Q., Wang, Z., Liu, Z., *et al.* (2022) A new troodontid from the Upper Cretaceous Gobi Basin of inner Mongolia, China. *Cretaceous Research*, **130**, 105052.

Pérez, M.E., Vallejo-Pareja, M.C., Carrillo, J.D. & Jaramillo, C. (2017) A New Pliocene Capybara (Rodentia, Caviidae) from Northern South America (Guajira, Colombia), and its Implications for the Great American Biotic Interchange. *Journal of Mammalian Evolution*, **24**, 111–125.

Peyre de Fabrègues, C., Bi, S., Li, H., Li, G., Yang, L. & Xu, X. (2020) A new species of early-diverging Sauropodiformes from the Lower Jurassic Fengjiahe Formation of Yunnan Province, China. *Scientific Reports*, **10**, 10961.

Pinheiro, F.L., Silva-Neves, E. & Da-Rosa, Á.A.S. (2021) An early-diverging procolophonid from the lowermost Triassic of South America and the origins of herbivory in Procolophonoidea. *Papers in Palaeontology*, **7**, 1601–1612.

Poole, K. (2022) Phylogeny of iguanodontian dinosaurs and the evolution of quadrupedality. *Palaeontologia Electronica*, **25**, 1–65.

Poust, A.W., Gao, C., Varricchio, D.J., Wu, J. & Zhang, F. (2020) A new microraptorine theropod from the Jehol Biota and growth in early dromaeosaurids. *The Anatomical Record*, **303**, 963–987.

Praschag, P., Schmidt, C., Fritzsch, G., Müller, A., Gemel, R. & Fritz, U. (2006) *Geoemyda silvatica*, an enigmatic turtle of the Geoemydidae (Reptilia: Testudines), represents a distinct genus. *Organisms Diversity & Evolution*, **6**, 151–162.

Prieto-Márquez, A. (2010) Global phylogeny of Hadrosauridae (Dinosauria: Ornithopoda) using parsimony and Bayesian methods. *Zoological Journal of the Linnean Society*, **159**, 435–502.

Pritchard, A.C. & Nesbitt, S.J. (2017) A bird-like skull in a Triassic diapsid reptile increases heterogeneity of the morphological and phylogenetic radiation of Diapsida. *Royal Society Open Science*, **4**, 170499.

Pritchard, A.C. & Sues, H.-D. (2019) Postcranial remains of *Teraterpeton hrynewichorum* (Reptilia: Archosauromorpha) and the mosaic evolution of the saurian postcranial skeleton. *Journal of Systematic Palaeontology*.

Pritchard, A.C., Sues, H.-D., Scott, D. & Reisz, R.R. (2021) Osteology, relationships and functional morphology of *Weigeltisaurus jaekeli* (Diapsida, Weigeltisauridae) based on a complete skeleton from the Upper Permian Kupferschiefer of Germany. *PeerJ*, **9**, e11413.

Pritchard, A.C., Turner, A.H., Irmis, R.B., Nesbitt, S.J. & Smith, N.D. (2016) Extreme Modification of the Tetrapod Forelimb in a Triassic Diapsid Reptile. *Current Biology*, **26**, 2779–2786.

Puértolas-Pascual, E., Canudo, J.I. & Moreno-Azanza, M. (2014) The eusuchian crocodylomorph *Allodaposuchus subjuniperus* sp. nov., a new species from the latest Cretaceous (upper Maastrichtian) of Spain. *Historical Biology*, **26**, 91–109.

Pyron, R.A. (2017) Novel Approaches for Phylogenetic Inference from Morphological Data and Total-Evidence Dating in Squamate Reptiles (Lizards, Snakes, and Amphisbaenians). *Systematic Biology*, **66**, 38–56.

Qin, Z., Clark, J., Choiniere, J. & Xu, X. (2019) A new alvarezsaurian theropod from the Upper Jurassic Shishugou Formation of western China. *Scientific Reports*, **9**, 11727.

Rauhut, O.W.M., Foth, C., Tischlinger, H. & Norell, M.A. (2012) Exceptionally preserved juvenile megalosauroid theropod dinosaur with filamentous integument from the Late Jurassic of Germany. *Proceedings of the National Academy of Sciences*, **109**, 11746–11751.

Rauhut, O.W.M., Holwerda, F.M. & Furrer, H. (2020) A derived sauropodiform dinosaur and other sauropodomorph material from the Late Triassic of Canton Schaffhausen, Switzerland. *Swiss Journal of Geosciences*, **113**, 8.

Rauhut, O.W.M., Hübner, T. & Lanser, K.-P. (2016) A new megalosaurid theropod dinosaur from the late Middle Jurassic (Callovian) of north-western Germany: implications for theropod evolution and faunal turnover in the Jurassic. *Palaeontologia Electronica*, **19**, 1–65.

Rauhut, O.W.M. & Pol, D. (2019) Probable basal allosauroid from the early Middle Jurassic Cañadón Asfalto Formation of Argentina highlights phylogenetic uncertainty in tetanuran theropod dinosaurs. *Scientific Reports*, **9**, 18826.

Raven, T.J., Barrett, P.M., Joyce, C.B. & Maidment, S.C.R. (2023) The phylogenetic relationships and evolutionary history of the armoured dinosaurs (Ornithischia: Thyreophora). *Journal of Systematic Palaeontology*, **21**, 2205433.

Raven, T.J. & Maidment, S.C.R. (2017) A new phylogeny of Stegosauria (Dinosauria, Ornithischia). *Palaeontology*, **60**, 401–408.

Reisz, R.R. & Fröbisch, J. (2014) The Oldest Caseid Synapsid from the Late Pennsylvanian of Kansas, and the Evolution of Herbivory in Terrestrial Vertebrates. *PLOS ONE*, **9**, e94518.

Ren, X.-X., Jiang, S., Wang, X.-R., Peng, G.-Z., Ye, Y., King, L., *et al.* (2022) Osteology of *Dashanpusaurus dongi* (Sauropoda: Macronaria) and new evolutionary evidence from Middle Jurassic Chinese sauropods. *Journal of Systematic Palaeontology*, **20**, 1–72.

Reyes, W.A., Parker, W.G. & Marsh, A.D. (2020) Cranial anatomy and dentition of the aetosaur *Typothorax coccinarum* (Archosauria: Pseudosuchia) from the Upper Triassic (Revueltian–mid Norian) Chinle Formation of Arizona. *Journal of Vertebrate Paleontology*, **40**, e1876080.

Rincón, A.F., Raad Pájaro, D.A., Jiménez Velandia, H.F., Ezcurra, M.D. & Wilson Mantilla, J.A. (2022) A sauropod from the Lower Jurassic La Quinta Formation (Dept. Cesar, Colombia) and the initial diversification of eusauropods at low latitudes. *Journal of Vertebrate Paleontology*, **42**, e2077112.

Rio, J.P. & Mannion, P.D. (2021) Phylogenetic analysis of a new morphological dataset elucidates the evolutionary history of Crocodylia and resolves the long-standing gharial problem. *PeerJ*, **9**, e12094.

Romano, M., Ronchi, A., Maganuco, S. & Nicosia, U. (2017) New material of *Alierasaurus ronchii* (Synapsida, Caseidae) from the Permian of Sardinia (Italy), and its phylogenetic affinities. *Palaeontologia Electronica*.

Romo De Vivar, P.R., Martinelli, A.G., Fonseca, P.H.M. & Soares, M.B. (2020) To be or not to be: The Hidden Side of *Cargninia Enigmatica* and Other Puzzling Remains of Lepidosauromorpha from the Upper Triassic of Brazil. *Journal of Vertebrate Paleontology*, **40**, e1828438.

Ruiz, P.G., Fuente, M.S. de la & Fernández, M.S. (2020) New cranial fossils of the Jurassic turtle *Neusticemys neuquina* and phylogenetic relationships of the only thalassochelydian known from the eastern Pacific. *Journal of Paleontology*, **94**, 145–164.

Ruta, M. & Bolt, J.R. (2008) The Brachyopoid *Hadrokkosaurus bradyi* from the Early Middle Triassic of Arizona, and a Phylogenetic Analysis of Lower Jaw Characters in Temnospondyl Amphibians. *Acta Palaeontologica Polonica*, **53**, 579–592.

Ruta, M., Clack, J.A. & Smithson, T.R. (2020) A review of the stem amniote *Eldeceeon rolfei* from the Viséan of East Kirkton, Scotland. *Earth and Environmental Science Transactions of The Royal Society of Edinburgh*, **111**, 173–192.

Ruta, M., Jeffery, J.E. & Coates, M.I. (2003) A supertree of early tetrapods. *Proceedings of the Royal Society of London. Series B: Biological Sciences*, **270**, 2507–2516.

Ruta, M., Krieger, J., Angielczyk, K.D. & Wills, M.A. (2018) The evolution of the tetrapod humerus: morphometrics, disparity, and evolutionary rates. *Earth and Environmental Science Transactions of The Royal Society of Edinburgh*, **109**, 351–369.

Ryan, M.J., Evans, D.C. & Shepherd, K.M. (2012) A new ceratopsid from the Foremost Formation (middle Campanian) of Alberta. *Canadian Journal of Earth Sciences*, **49**, 1251–1262.

Sánchez-Villagra, M.R., Kay, R.F. & Anaya-Daza, F. (2000) Cranial anatomy and palaeobiology of the Miocene marsupial *Hondalagus altiplanensis* and a phylogeny of argyrolagids. *Palaeontology*, **43**, 287–301.

Sander, P.M. (1989) The pachypleurosaurids (Reptilia: Nothosauria) from the Middle Triassic of Monte San Giorgio (Switzerland) with the description of a new species. *Philosophical Transactions of the Royal Society of London. B, Biological Sciences*, **325**, 561–666.

Sasso, C.D., Maganuco, S. & Cau, A. (2018) The oldest ceratosaurian (Dinosauria: Theropoda), from the Lower Jurassic of Italy, sheds light on the evolution of the three-fingered hand of birds. *PeerJ*, **6**, e5976.

Scavezzoni, I. & Fischer, V. (2018) *Rhinochelys amaberti* Moret (1935), a protostegid turtle from the Early Cretaceous of France. *PeerJ*, **6**, e4594.

Schoch, R.R. (2012) Character distribution and phylogeny of the dissorophid temnospondyls. *Fossil Record*, **15**, 121–137.

Schoch, R.R. (2013) The evolution of major temnospondyl clades: an inclusive phylogenetic analysis. *Journal of Systematic Palaeontology*, **11**, 673–705.

Schoch, R.R. (2018a) Osteology of the temnospondyl *Neldasaurus* and the evolution of basal dvinosaurians. *Neues Jahrbuch für Geologie und Paläontologie - Abhandlungen*, 1–16.

Schoch, R.R. (2018b) The temnospondyl *Parotosuchus nasutus* (v. Meyer, 1858) from the Early Triassic Middle Buntsandstein of Germany. *Palaeodiversity*, **11**, 107–126.

Schoch, R.R. (2021) Osteology of the Permian temnospondyl amphibian *Glanochthon lellbachae* and its relationships. *Fossil Record*, **24**, 49–64.

Schoch, R.R. & Milner, A.R. (2008) The intrarelationships and evolutionary history of the temnospondyl family Branchiosauridae. *Journal of Systematic Palaeontology*, **6**, 409–431.

Schoch, R.R., Werneburg, R. & Voigt, S. (2020) A Triassic stem-salamander from Kyrgyzstan and the origin of salamanders. *Proceedings of the National Academy of Sciences*, **117**, 11584–11588.

Schoch, R.R. & Witzmann, F. (2009) Osteology and relationships of the temnospondyl genus *Sclerocephalus*. *Zoological Journal of the Linnean Society*, **157**, 135–168.

Schulp, A.S., Jagt, J.W.M. & Fonken, F. (2004) New material of the mosasaur *Carinodens belgicus* from the Upper Cretaceous of the Netherlands. *Journal of Vertebrate Paleontology*, **24**, 744–747.

Sequeira, S.E.K. (2003) The skull of *Cochleosaurus bohemicus* Frič, a temnospondyl from the Czech Republic (Upper Carboniferous) and cochleosaurid interrelationships. *Earth and Environmental Science Transactions of The Royal Society of Edinburgh*, **94**, 21–43.

Shen, C., Lü, J., Liu, S., Kundrát, M., Brusatte, S.L. & Gao, H. (2017a) A New Troodontid Dinosaur from the Lower Cretaceous Yixian Formation of Liaoning Province, China. *Acta Geologica Sinica - English Edition*, **91**, 763–780.

Shen, C., Zhao, B., Gao, C., Lü, J. & Kundrát, M. (2017b) A new troodontid dinosaur (*Liaoningvenator curriei* gen. et sp. nov.) from the Early Cretaceous Yixian Formation of Liaoning Province. *Acta Geologica Sinica - English Edition*, **38**, 359–371.

Sidor, C.A., Kulik, Z.T. & Huttenlocker, A.K. (2022) A new bauriamorph therocephalian adds a novel component to the Lower Triassic tetrapod assemblage of the Fremouw Formation (Transantarctic Basin) of Antarctica. *Journal of Vertebrate Paleontology*, **41**, e2081510.

Sigurdsen, T. & Bolt, J.R. (2010) The Lower Permian amphibamid *Doleserpeton* (Temnospondyli: Dissorophoidea), the interrelationships of amphibamids, and the origin of modern amphibians. *Journal of Vertebrate Paleontology*, **30**, 1360–1377.

Simões, T.R., Caldwell, M.W. & Kellner, A.W.A. (2015a) A new Early Cretaceous lizard species from Brazil, and the phylogenetic position of the oldest known South American squamates. *Journal of Systematic Palaeontology*, **13**, 601–614.

Simões, T.R., Caldwell, M.W. & Pierce, S.E. (2020) Sphenodontian phylogeny and the impact of model choice in Bayesian morphological clock estimates of divergence times and evolutionary rates. *BMC Biology*, **18**, 191.

Simões, T.R., Caldwell, M.W., Tałanda, M., Bernardi, M., Palci, A., Vernygora, O., *et al.* (2018) The origin of squamates revealed by a Middle Triassic lizard from the Italian Alps. *Nature*, **557**, 706–709.

Simões, T.R., Vernygora, O., Paparella, I., Jimenez-Huidobro, P. & Caldwell, M.W. (2017) Mosasauroid phylogeny under multiple phylogenetic methods provides new insights on the evolution of aquatic adaptations in the group. *PLOS ONE*, **12**, e0176773.

Simões, T.R., Wilner, E., Caldwell, M.W., Weinschütz, L.C. & Kellner, A.W.A. (2015b) A stem acrodontan lizard in the Cretaceous of Brazil revises early lizard evolution in Gondwana. *Nature Communications*, **6**, 8149.

Skutschas, P.P. & Gubin, Y.M. (2012) A New Salamander from the Late Paleocene—Early Eocene of Ukraine. *Acta Palaeontologica Polonica*, **57**, 135–148.

Smith, K.T. & Gauthier, J.A. (2013) Early Eocene Lizards of the Wasatch Formation near Bitter Creek, Wyoming: Diversity and Paleoenvironment during an Interval of Global Warming. *Bulletin of the Peabody Museum of Natural History*, **54**, 135–230.

Solé, F. (2013) New proviverrine genus from the Early Eocene of Europe and the first phylogeny of Late Palaeocene–Middle Eocene hyaenodontidans (Mammalia). *Journal of Systematic Palaeontology*, **11**, 375–398.

Sookias, R.B. (2016) The relationships of the Euparkeriidae and the rise of Archosauria. *Royal Society Open Science*, **3**, 150674.

Souza, G.A. de, Soares, M.B., Weinschütz, L.C., Wilner, E., Lopes, R.T., Araújo, O.M.O. de, *et al.* (2021) The first edentulous ceratosaur from South America. *Scientific Reports*, **11**, 22281.

Spindler, F. (2015) *The basal Spenacodontia - systematic revision and evolutionary implications* (PhD Thesis).

Steeman, M.E. (2007) Cladistic analysis and a revised classification of fossil and recent mysticetes. *Zoological Journal of the Linnean Society*, **150**, 875–894.

Sterli, J. & De La Fuente, M.S. (2011) Re-description and evolutionary remarks on the Patagonian horned turtle *Niolamia argentina* Ameghino, 1899 (Testudinata, Meiolaniidae). *Journal of Vertebrate Paleontology*, **31**, 1210–1229.

Sterli, J., Fuente, M.S. de la & Rougier, G.W. (2018) New remains of *Condorchelys antiqua* (Testudinata) from the Early-Middle Jurassic of Patagonia: anatomy, phylogeny, and paedomorphosis in the early evolution of turtles. *Journal of Vertebrate Paleontology*.

Suchkova, J. & Golubev, V. (2019) A New Primitive Therocephalian (Theromorpha) from the Middle Permian of Eastern Europe. *Paleontological Journal*, **53**, 88–96.

Sues, H.-D., Fitch, A.J. & Whatley, R.L. (2020) A New Rhynchosaur (Reptilia, Archosauromorpha) from the Upper Triassic of Eastern North America. *Journal of Vertebrate Paleontology*, **40**, e1771568.

Sues, H.-D. & Kligman, B.T. (2021) A new lizard-like reptile from the Upper Triassic (Carnian) of Virginia and the Triassic record of Lepidosauromorpha (Diapsida, Sauria). *Journal of Vertebrate Paleontology*, **40**, e1879102.

Sues, H.-D. & Schoch, R.R. (2013) Reassessment of cf. *Halticosaurus orbitoangulatus* from the Upper Triassic (Norian) of Germany–a pseudosuchian, not a dinosaur. *Zoological Journal of the Linnean Society*, **168**, 859–872.

Swartz, B. (2012) A Marine Stem-Tetrapod from the Devonian of Western North America. *PLOS ONE*, **7**, e33683.

Sweetman, S.C. (2004) The first record of velociraptorine dinosaurs (Saurischia, Theropoda) from the Wealden (Early Cretaceous, Barremian) of southern England. *Cretaceous Research*, **25**, 353–364.

Tałanda, M., Fernandez, V., Panciroli, E., Evans, S.E. & Benson, R.J. (2022) Synchrotron tomography of a stem lizard elucidates early squamate anatomy. *Nature*, **611**, 99–104.

Tambussi, C., Degrange, F.J., De Mendoza, R.S., Sferco, E. & Santillana, S. (2019) A stem anseriform from the early Palaeocene of Antarctica provides new key evidence in the early evolution of waterfowl.pdf. *Zoological Journal of the Linnean Society*, **XX**, 1–28.

Tolchard, F., Smith, R.M.H., Arcucci, A., Mocke, H. & Choiniere, J.N. (2021) A new 'rauisuchian' archosaur from the Middle Triassic Omingonde Formation (Karoo Supergroup) of Namibia. *Journal of Systematic Palaeontology*, **19**, 595–631.

Tong, H., Claude, J., Li, C.-S., Yang, J. & Smith, T. (2019) *Wutuchelys eocenica* n. gen. n. sp., an Eocene stem testudinoid turtle from Wutu, Shandong Province, China. *Geological Magazine*, **156**, 133–146.

Tschopp, E. & Mateus, O. (2017) Osteology of *Galeamopus pabsti* sp. nov. (Sauropoda: Diplodocidae), with implications for neurocentral closure timing, and the cervico-dorsal transition in diplodocids. *PeerJ*, **5**, e3179.

Tsuji, L.A. & Müller, J. (2009) Assembling the history of the Parareptilia: phylogeny, diversification, and a new definition of the clade. *Fossil Record*, **12**, 71–81.

Tsuji, L.A., Müller, J. & Reisz, R.R. (2010) *Microleter mckinzieorum* gen. et sp. nov. from the Lower Permian of Oklahoma: the basalmost parareptile from Laurasia. *Journal of Systematic Palaeontology*, **8**, 245–255.

Tsuji, L.A., Sobral, G. & Müller, J. (2013) *Ruhuhuaria reiszi*, a new procolophonoid reptile from the Triassic Ruhuhu Basin of Tanzania. *Comptes Rendus Palevol*, A tribute to Robert R. Reisz / Un hommage à Robert R. Reisz, **12**, 487–494.

Upchurch, P., Barrett, P.M., Xijin, Z. & Xing, X. (2007) A re-evaluation of *Chinshakiangosaurus chunghoensis* Ye *vide* Dong 1992 (Dinosauria, Sauropodomorpha): implications for cranial evolution in basal sauropod dinosaurs. *Geological Magazine*, **144**, 247–262.

Upham, N.S., Esselstyn, J.A. & Jetz, W. (2019) Inferring the mammal tree: Species-level sets of phylogenies for questions in ecology, evolution, and conservation. *PLOS Biology*, **17**, e3000494.

Utzeri, V.J., Cilli, E., Fontani, F., Zoboli, D., Orsini, M., Ribani, A., *et al.* (2023) Ancient DNA re-opens the question of the phylogenetic position of the Sardinian pika *Prolagus sardus* (Wagner, 1829), an extinct lagomorph. *Scientific Reports*, **13**, 13635.

Vallin, G. & Laurin, M. (2004) Cranial morphology and affinities of *Microbrachis* , and a reappraisal of the phylogeny and lifestyle of the first amphibians. *Journal of Vertebrate Paleontology*, **24**, 56–72.

Vasilyev, V.A., Korsunenko, A.V., Pereshkolnik, S.L., Mazanaeva, L.F., Bannikova, A.A., Bondarenko, D.A., *et al.* (2014) Differentiation of tortoises of the genera *Testudo* and *Agrionemys* (Testudinidae) based on the polymorphism of nuclear and mitochondrial markers. *Russian Journal of Genetics*, **50**, 1060–1074.

Velazco, P.M., Buczek, A.J. & Novacek, M.J. (2017) Two New Tritylodontids (Synapsida, Cynodontia, Mammaliamorpha) from the Upper Jurassic, Southwestern Mongolia. *American Museum Novitates*, **2017**, 1–35.

Venczel, M. & Codrea, V.A. (2018) A new proteid salamander from the early Oligocene of Romania with notes on the paleobiogeography of Eurasian proteids. *Journal of Vertebrate Paleontology*, **38**, e1508027.

Villa, A., Abella, J., Alba, D.M., Almécija, S., Bolet, A., Koufos, G.D., *et al.* (2018) Revision of *Varanus marathonensis* (Squamata, Varanidae) based on historical and new material: morphology, systematics, and paleobiogeography of the European monitor lizards. *PLOS ONE*, **13**, e0207719.

Villa, A., Wings, O. & Rabi, M. (2022) A new gecko (Squamata, Gekkota) from the Eocene of Geiseltal (Germany) implies long-term persistence of European Sphaerodactylidae. *Papers in Palaeontology*, **8**, e1434.

Vincent, P. & Storrs, G.W. (2019) *Lindwurmia*, a new genus of Plesiosauria (Reptilia: Sauropterygia) from the earliest Jurassic of Halberstadt, northwest Germany. *The Science of Nature*, **106**, 5.

Voris, J.T., Therrien, F., Zelenitsky, D.K. & Brown, C.M. (2020) A new tyrannosaurine (Theropoda:Tyrannosauridae) from the Campanian Foremost Formation of Alberta, Canada, provides insight into the evolution and biogeography of tyrannosaurids. *Cretaceous Research*, **110**, 104388.

Wang, W., Ma, F. & Li, C. (2020a) First subadult specimen of Psephochelys polyosteoderma (Sauropterygia, Placodontia) implies turtle-like fusion pattern of the carapace. *Papers in Palaeontology*, **6**, 251–264.

Wang, X., Huang, J., Kundrát, M., Cau, A., Liu, X., Wang, Y., *et al.* (2020b) A new jeholornithiform exhibits the earliest appearance of the fused sternum and pelvis in the evolution of avialan dinosaurs. *Journal of Asian Earth Sciences*, **199**, 104401.

Wang, Y. (2004) Taxonomy and Stratigraphy of Late Mesozoic Anurans and Urodeles from China. *Acta Geologica Sinica - English Edition*, **78**, 1169–1178.

Warren, A. & Marsicano, C. (2000) A phylogeny of the Brachyopoidea (Temnospondyli, Stereospondyli). *Journal of Vertebrate Paleontology*, **20**, 462–483.

Wencker, L.C.M., Tschopp, E., Villa, A., Augé, M.L. & Delfino, M. (2021) Phylogenetic value of jaw elements of lacertid lizards (Squamata: Lacertoidea): a case study with Oligocene material from France. *Cladistics*, **37**, 765–802.

Werneburg, R., Spindler, F., Falconnet, J., Steyer, J.-S., liaud, M. vianey- & Schneider, J. (2022) A new caseid synapsid from the Permian (Guadalupian) of the Lodève basin (Occitanie, France), **45**, 1–36.

Whitney, M.R. & Sidor, C.A. (2016) A new therapsid from the Permian Madumabisa Mudstone Formation (Mid-Zambezi Basin) of southern Zambia. *Journal of Vertebrate Paleontology*, **36**, e1150767.

Whitney, M.R. & Sidor, C.A. (2019) Histological and developmental insights into the herbivorous dentition of tapinocephalid therapsids. *PLOS ONE*, **14**, e0223860.

Wible, J.R., Novacek, M.J. & Rougier, G.W. (2004) New data on the skull and dentition in the Mongolian Late Cretaceous eutherian mammal *Zalambdalestes*. *Bulletin of the American Museum of Natural History*, **2004**, 1–144.

Wible, J.R., Rougier, G.W., Novacek, M.J. & Asher, R.J. (2007) Cretaceous eutherians and Laurasian origin for placental mammals near the K/T boundary. *Nature*, **447**, 1003–1006.

Wible, J.R., Rougier, G.W., Novacek, M.J. & Asher, R.J. (2009) The Eutherian Mammal *Maelestes gobiensis* from the Late Cretaceous of Mongolia and the phylogeny of Cretaceous Eutheria. *Bulletin of the American Museum of Natural History*, **2009**, 1–123.

Wiens, J.J., Kuczynski, C.A., Townsend, T., Reeder, T.W., Mulcahy, D.G. & Sites, J.W., Jr. (2010) Combining Phylogenomics and Fossils in Higher-Level Squamate Reptile Phylogeny: Molecular Data Change the Placement of Fossil Taxa. *Systematic Biology*, **59**, 674–688.

Wilberg, E.W., Turner, A.H. & Brochu, C.A. (2019) Evolutionary structure and timing of major habitat shifts in Crocodylomorpha. *Scientific Reports*, **9**, 514.

Windholz, G.J., Coria, R.A., Bellardini, F., Baiano, M.A., Pino, D.A., Ortega, F., *et al.* (2022) On a dicraeosaurid specimen from the Mulichinco Formation (Valanginian, Neuquén Basin) of Argentina and phylogenetic relationships of the South American dicraeosaurids (Sauropoda, Diplodocoidea).

Witzmann, F. & Ruta, M. (2018) Evolutionary changes in the orbits and palatal openings of early tetrapods, with emphasis on temnospondyls. *Earth and Environmental Science Transactions of the Royal Society of Edinburgh*, **109**, 333–350.

Witzmann, F. & Schoch, R.R. (2018) Skull and postcranium of the bystrowianid *Bystrowiella schumanni* from the Middle Triassic of Germany, and the position of chroniosuchians within Tetrapoda. *Journal of Systematic Palaeontology*, **16**, 711–739.

Wolniewicz, A.S., Shen, Y., Li, Q., Sun, Y., Qiao, Y., Chen, Y., *et al.* (2023) An armoured marine reptile from the Early Triassic of South China and its phylogenetic and evolutionary implications. *eLife*, **12**, e83163.

Woodruff, D.C., Goodwin, M.B., Lyson, T.R. & Evans, D.C. (2021) Ontogeny and variation of the pachycephalosaurine dinosaur Sphaerotholus buchholtzae, and its systematics within the genus. *Zoological Journal of the Linnean Society*, **193**, 563–601.

Woods, R., Barnes, I., Brace, S. & Turvey, S.T. (2021) Ancient DNA Suggests Single Colonization and Within-Archipelago Diversification of Caribbean Caviomorph Rodents. *Molecular Biology and Evolution*, **38**, 84–95.

Worthy, T.H., Degrange, F.J., Handley, W.D. & Lee, M.S.Y. (2017) The evolution of giant flightless birds and novel phylogenetic relationships for extinct fowl (Aves, Galloanseres). *Royal Society Open Science*, **4**, 170975.

Wu, X.-C., Shi, J.-R., Dong, L.-Y., Carr, T.D., Yi, J. & Xu, S.-C. (2020) A new tyrannosauroid from the Upper Cretaceous of Shanxi, China. *Cretaceous Research*, **108**, 104357.

Xing, L., Stanley, E.L., Bai, M. & Blackburn, D.C. (2018) The earliest direct evidence of frogs in wet tropical forests from Cretaceous Burmese amber. *Scientific Reports*, **8**, 8770.

Xu, G.-H., Ren, Y., Zhao, L.-J., Liao, J.-L. & Feng, D.-H. (2022a) A long-tailed marine reptile from China provides new insights into the Middle Triassic pachypleurosaur radiation. *Scientific Reports*, **12**, 7396.

Xu, L., Wu, X., Junchang, L., Jia, S., Zhang, J., Pu, H., *et al.* (2014) A New Lizard (Lepidosauria: Squamata) from the Upper Cretaceous of Henan, China. *Acta Geologica Sinica - English Edition*, **88**, 1041–1050.

Xu, X., Zhao, X. & Clark, J.M. (2001) A new therizinosaur from the Lower Jurassic lower Lufeng Formation of Yunnan, China. *Journal of Vertebrate Paleontology*, **21**, 477–483.

Xu, Y., Jiang, S. & Wang, X. (2022b) A new istiodactylid pterosaur, *Lingyuanopterus camposi* gen. et sp. nov., from the Jiufotang Formation of western Liaoning, China. *PeerJ*, **10**, e13819.

Yang, Y., Wu, W., Dieudonné, P.-E. & Godefroit, P. (2020) A new basal ornithopod dinosaur from the Lower Cretaceous of China. *PeerJ*, **8**, e9832.

Yates, A.M. & Sengupta, D.P. (2002) A lapillopsid temnospondyl from the Early Triassic of India. *Alcheringa: An Australasian Journal of Palaeontology*, **26**, 201–208.

Young, M.T., Brusatte, S., Ruta, M. & De Andrade, M.B. (2010) The evolution of Metriorhynchoidea (mesoeucrocodylia, thalattosuchia): an integrated approach using geometric morphometrics, analysis of disparity, and biomechanics. *Zoological Journal of the Linnean Society*, **158**, 801–859.

Yu, C., Prieto-Marquez, A., Chinzorig, T., Badamkhatan, Z. & Norell, M. (2020) A neoceratopsian dinosaur from the early Cretaceous of Mongolia and the early evolution of ceratopsia. *Communications Biology*, **3**, 1–8.

Zaher, H., Pol, D., Navarro, B.A., Delcourt, R. & Carvalho, A.B. (2020) An Early Cretaceous theropod dinosaur from Brazil sheds light on the cranial evolution of the Abelisauridae. *Comptes Rendus Palevol*.

Zanno, L.E. (2010) A taxonomic and phylogenetic re-evaluation of Therizinosauria (Dinosauria: Maniraptora). *Journal of Systematic Palaeontology*, **8**, 503–543.

Zhou, C.-F., Gao, K.-Q., Yi, H., Xue, J., Li, Q. & Fox, R.C. (2017) Earliest filter-feeding pterosaur from the Jurassic of China and ecological evolution of Pterodactyloidea. *Royal Society Open Science*, **4**, 160672.

Zhou, C.-F., Rabi, M. & Joyce, W.G. (2014) A new specimen of Manchurochelys manchoukuoensis from the Early Cretaceous Jehol Biota of Chifeng, Inner Mongolia, China and the phylogeny of Cretaceous basal eucryptodiran turtles. *BMC Evolutionary Biology*, **14**, 77.

Zhu, M., Ahlberg, P.E., Zhao, W.-J. & Jia, L.-T. (2017) A Devonian tetrapod-like fish reveals substantial parallelism in stem tetrapod evolution. *Nature Ecology & Evolution*, **1**, 1470–1476.

Zrzavý, J., Duda, P., Robovský, J., Okřinová, I. & Pavelková Říčánková, V. (2018) Phylogeny of the Caninae (Carnivora): Combining morphology, behaviour, genes and fossils. *Zoologica Scripta*, **47**, 373–389.
